# An energy information trade-off explains hidden structure in the neuronal parameter space

**DOI:** 10.64898/2026.03.02.708507

**Authors:** Philip Sommer, Fleur Zeldenrust, Peter Jedlicka, Alexander D. Bird, Jochen Triesch

**Affiliations:** Frankfurt Institute for Advanced Studies, Frankfurt am Main, Germany; Department of Physiology, Anatomy and Genetics, University of Oxford, Oxford, United Kingdom; Donders Institute for Brain, Behaviour and Cognition, Radboud University, Nijmegen, The Netherlands; Computer-Based Modelling in the field of 3R Animal Protection, ICAR3R, Faculty of Medicine, Justus Liebig University, Giessen, Germany; Translational Neuroscience Network Giessen, Germany; Institute of Clinical Neuroanatomy, Neuroscience Center, Goethe University, Frankfurt; Ernst Strüngmann Institute (ESI) for Neuroscience in cooperation with the Max Planck Society, Frankfurt am Main, Germany

**Keywords:** neuronal diversity, neuronal energy budget, energy-efficiency, Pareto-optimality, degeneracy, food restriction, synaptic scaling

## Abstract

Among neurons of the same type, the different electrophysiological parameters vary drastically. It is presently unclear if there is any hidden structure in this diversity and how it relates to neuronal function. One potential source of structure is the evolutionary pressure that has sculpted brains to work in an energy-efficient manner. Here we show that neurons’ parameters do not vary randomly, but that neurons populate a manifold within their parameter space permitting energy-efficient signaling. This manifold represents a Pareto front of degenerate solutions to the problem of energy-efficiency and explains the prevalence of low to intermediate firing rates (2 – 5 Hz) in neocortex. Furthermore, we show that neurons in different sensory brain areas populate different manifolds and that food restriction induces systematic shifts of neuronal parameters along this manifold. Our results suggest that the large diversity of neuronal properties is actually tightly regulated to ensure energy-efficient signaling in different contexts.

**In brief:** We show that there is hidden structure in the diversity of electrophysiological properties of neurons that is governed by trade-offs between energy consumption and information transmission.

**Highlights:**

- We derive an updated neuronal energy budget for a large class of neuron models.
- Computational modeling predicts a manifold of degenerate solutions to energy-efficient information coding forming a Pareto front in the neuronal parameter space.
- Neurons from visual and somatosensory cortex fall on the predicted manifold.
- Optimal energy-efficiency is achieved for intermediate-low firing rates (2 – 5 Hz) as observed in neocortex.
- We observe an energy-efficient form of non-multiplicative synaptic scaling during food deprivation.

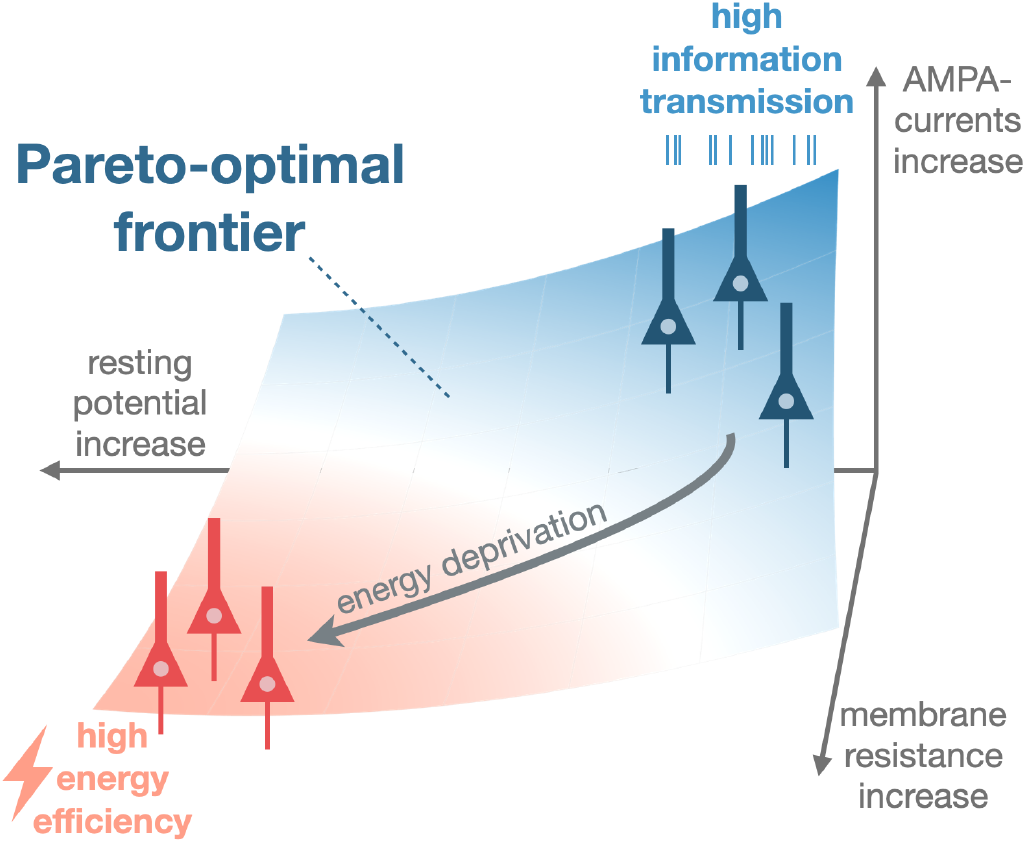

## Introduction

Neurons exhibit a striking variability in their intrinsic biophysical properties. Several studies (Marder and Goaillard, 2006; Goaillard and Marder, 2021) have investigated the vast space of neural parameters and shown that features such as ion channel densities can vary by two to three orders of magnitude across neurons of the same type. Work by Prinz *et al*. (2003, 2004) and Deistler et al. (2022) demonstrated that disparate, but well-balanced, combinations of synaptic and intrinsic parameters can give rise to similar functional outcomes (realistic neuronal and network behavior) (Marder and Goaillard, 2006), a phenomenon known as degeneracy. Degenerate solutions likely provide an evolutionary advantage by reducing the need for tight (homeostatic) regulation of individual ion channel levels while maintaining overall neuronal function (Marder and Prinz, 2002; Mishra and Narayanan, 2021; Stöber *et al*., 2023). Furthermore, heterogeneous response properties can enhance the quality of stimulus representation and coding range (Padmanabhan and Urban, 2010; Mejias and Longtin, 2012; Perez-Nieves *et al*., 2021; Gast *et al*., 2024; Dahmen *et al*., 2025). This flexibility also enables neurons to compensate for perturbations, such as ion channel knockouts (O’Leary *et al*., 2013), increasing robustness, adaptability, and potentially evolvability (Schneider et al., 2023). The question remains, however, how much of this variability comes from unavoidable, or unimportant, stochasticity in ion channel expression (Yang and Prescott, 2023) and how much is constrained by functional objectives.

Brains are energy-hungry, consuming 20% of the body’s total energy despite accounting for only 2% of human body mass (Wang et al., 2010) and evolution has driven them to work in an energy-efficient manner (Niven and Laughlin,2008, Hyder *et al*. 2013) . Energy availability is a constraint on brain size (Isler and Schaik, 2009; Navarrete *et al*., 2011), wiring patterns (Chen *et al*., 2006), synaptic transmission (Harris *et al*., 2015, 2019), neural coding (Levy and Baxter, 1996), and information processing (Schreiber et al., 2002) with multiple factors contributing to energy efficiency (Yu and Yu, 2017; Padamsey and Rochefort, 2023). In food restriction experiments, Padamsey et al. (2022) have recently observed that individual neurons in mouse visual cortex can reduce their energy consumption at the cost of information transmission. Specifically, these neurons appear to reduce energy consumption by changing their integration properties through the adaptation of membrane resistance, resting potential, and synaptic efficacies.

Trade-offs between functional performance, energy efficiency, robustness, and flexibility are ubiquitous in neural systems and impose fundamental constraints on neuronal design (Giudice and Crespi, 2018). A powerful framework for analyzing such multi-objective trade-offs is Pareto-optimality (Shoval *et al*., 2012; Remme *et al*., 2018; Pallasdies *et al*., 2021; Jedlicka *et al*., 2022), which describes optimal solutions balancing multiple competing objectives. A neuron is said to be Pareto optimal if its performance in one task, for example information coding, cannot be improved without reducing its performance in at least one other task, such as energy consumption. In this framework, alternative neuronal parameter sets should form a Pareto front with different points representing different trade-offs between the competing objectives.

We propose that a large part of experimentally observed neuronal heterogeneity can be understood as a manifestation of an information-energy trade-off, where neuronal phenotypes evolve towards and along a Pareto front maximizing information transmission while minimizing metabolic costs. Depending on the type of information encoded, availability of energy, and demanded precision, the energy-information trade-off might vary. To test this hypothesis, we analyze experimental data from two different primary sensory areas in rodents, each employing distinct coding strategies. We predict that the energy-information trade-off is a fundamental organizing principle of neural activity, shaping the observed diversity in neuronal properties.

Here we investigate energy-information trade-offs theoretically by combining data-driven simulations of an integrate-and-fire neuron model (Brette and Gerstner, 2005) and a detailed compartmental model (Goetz *et al*., 2021) with information measures (Shannon, 1948; Schreiber, 2000) and updated energy calculations (Attwell and Laughlin, 2001; Howarth *et al*., 2012; Yu *et al*., 2017). Our energy estimates take a new approach, being based on both first principles and ion-counting in biophysical models. Our work elucidates how the interplay between energy constraints and information transmission shapes neuronal diversity. We provide novel evidence that neuronal variability is not random, but resides in a constrained, low-dimensional subspace; a subspace that differs depending on coding task and energy availability. By demonstrating that neurons operate on a structured manifold of energy-efficient solutions, we provide a new perspective on degeneracy in neural systems, with neuronal parameter variability emerging from fundamental trade-offs rather than arbitrary fluctuations (Edelman and Gally, 2001). This framework offers a novel explanation for the observed variability in neuronal parameters and suggests that metabolic constraints play a key role in sculpting cortical firing rate distributions and coding strategies.

## Results

### Degenerate neuronal parameters reflect an optimally balanced trade-off between energy efficiency and information transmission

Neuronal parameters vary widely across brain regions and cell types in experimental recordings (Padamsey *et al*., 2022; Zeldenrust et al., 2024; Joshi et al., 2025) (Fig. 1A and 2A,B). We searched for structure in this variability by calculating the energy consumption of an adaptive exponential integrate-and-fire model (AdExp) (Brette and Gerstner, 2005) using our updated energy budget (see Methods). For visual cortex (Fig. 1) we systematically varied membrane resistance, resting potential, and synaptic weights — parameters experimentally shown to change under metabolic stress (Padamsey et al., 2022) — and evaluated neuronal performance in an orientation selectivity task (Fig. S1 — see Methods). For somatosensory cortex (Fig. 2) we explored the full parameter space extracted from experimental data (*V*_rest_, *R*_*m*_, *C*_*m*_, *V*_th_, *V*_reset_, Δ_*T*_, *τ*_ad_, *a*_ad_, *b*_ad_) (Zeldenrust *et al*., 2024) and assessed coding in a binary switching task (Fig. S11 — see Methods). We show that, rather than being arbitrary, the variability of neuronal parameters reflects the information-energy trade-offs made by different neurons in different situations. Using our newly derived estimates of neuronal energy consumption (see Methods), we identify a distinct subregion in the space of neuronal parameters where neurons transmit information in the most energy-efficient manner; this region lies close to the border where cells fall silent (Fig. 1B, shown as transparent circles). Experimental data from Padamsey *et al*. (2022) populate the predicted manifold (Fig. 1C), meaning that the recorded neurons lie along a structured Pareto-front representing optimal trade-offs between the two competing objectives of minimising energy consumption and maximising information transmission (proxied by orientation selectivity index, OSI). The Pareto front represents degenerate solutions to the biologically relevant tasks of the neuron.

**Figure 1.**
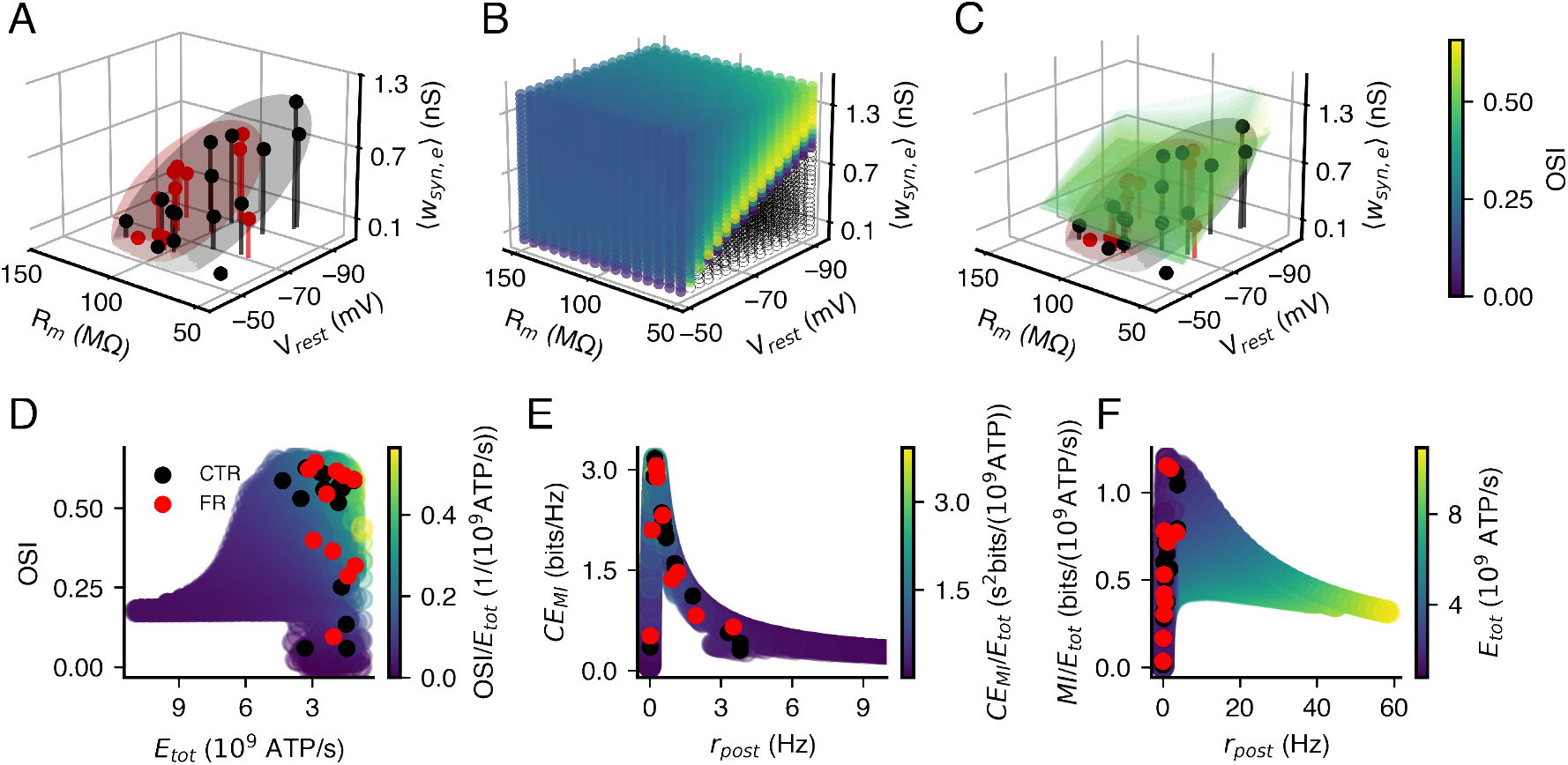
Visual cortex neurons balance energy efficiency and information transmission along a structured degeneracy manifold. **A**, Experimental data from food-restricted (FR, red) and control (CTR, black) animals (Padamse *al*., 2022) show both a high variability in neuronal parameters and systematic shifts under metabolic stress. FR neurons exhibit an increased membrane resistance, depolarized resting potentials, and reduced synaptic currents. Ellipsoids represent two standard deviations. **B**, Simulated parameter space (AdExp model with an orientation selectivity task (Fig. S1) — see Methods). Silent neurons are shown as transparent circles, while spiking neurons are color-coded by orientation selectivity index (OSI; colorbar in C). A structured subspace of high information transmission per unit energy emerges forming a Pareto-optimal manifold located near the boundary between active and silent regimes. **C**, Experimental data align with this predicted manifold, suggesting neurons maintain near-optimal trade-offs energy expenditure and information transmission. **D**, Visual cortex neurons (CTR and FR populate a Pareto front between energy efficiency and orientation selectivity (OSI). Color indicates OSI per expended energy. Total energy consumption (*E*_*tot*_) is computed using our updated energy budget (see Methods). **E**, Coding efficiency (mutual information per spike) is maximal at low-to-medium firing rates. Experimental neurons operate within this optimal range. Color indicates coding efficiency per unit energy. **F**, Mutual information per energy also peaks at low firing rates, offering an explanation for sparse cortical activity. Experimental cells prioritize information transmission over energy-efficiency thus populating the small *MI/E*_*tot*_ ratios left of the peak. Color indicates total energy expenditure.

**Figure 2.**
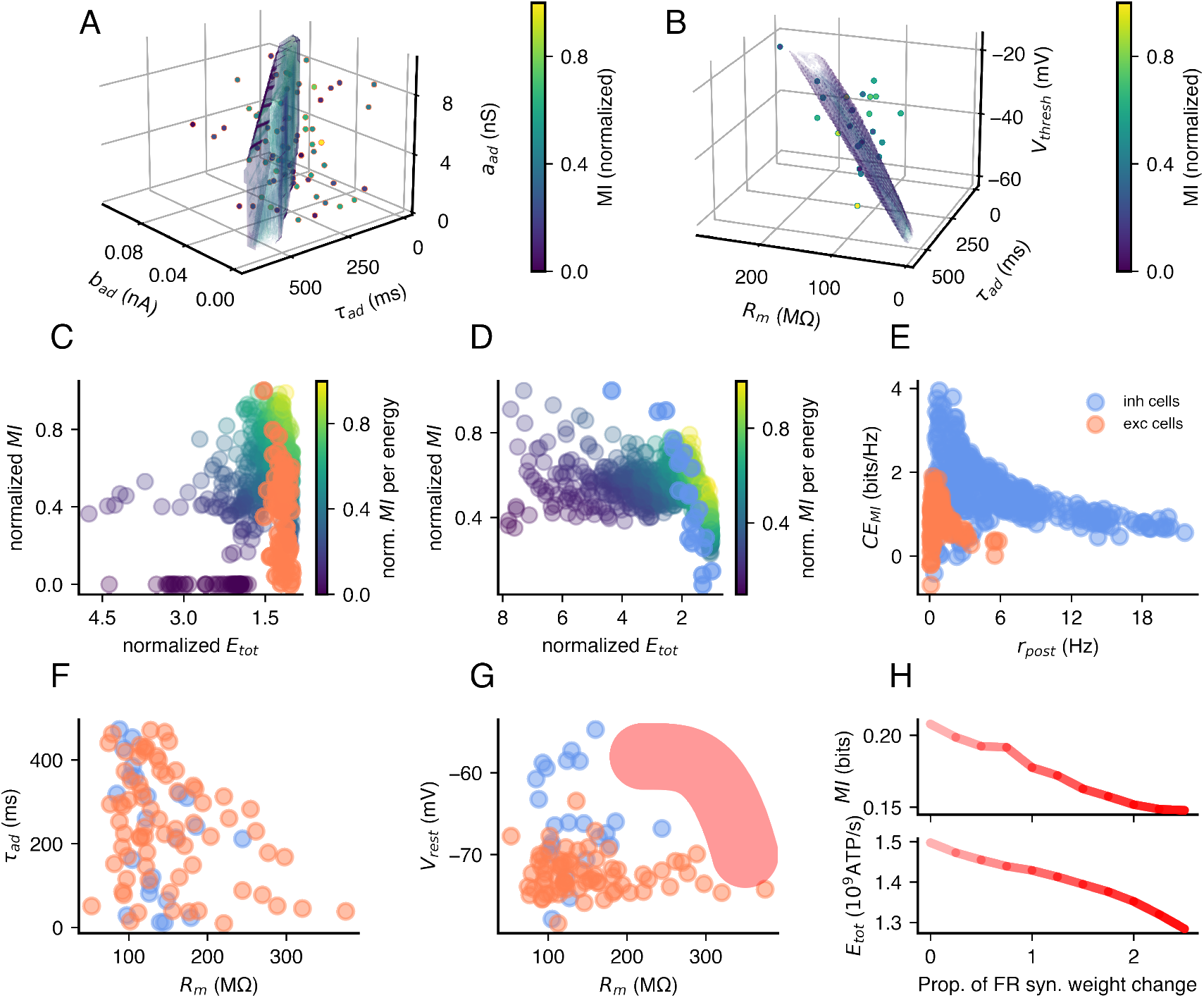
Distinct cortical areas and neuron types prioritize different energy–information trade offs. **A**, Excitatory and **B**, inhibitory barrel cortex neurons (Zeldenrust *et al*., 2024) shown as colored dots across three out of nine electrophysiological dimensions (*V*_rest_, *R*_*m*_, *C*_*m*_, *V*_th_, *V*_reset_, Δ_*T*_, *τ*_ad_, *a*_ad_, *b*_ad_ Simulated neurons (AdExp model with binary switching task (Fig. S11) — see Methods — parameters interpolated from 9D principal component space) shown as a semi-transparent volume colored by normalized mutual information (MI). **C**, Excitatory (orange) and **D**, inhibitory (light blue) neurons lie along distinct Pareto fronts spanning high energy efficiency and high MI (normalized to 1), reflecting cell-type-specific optimization strategies. **E**, Coding efficiency (MI/spike) differs in peak and shape between cell types, indicating different optimal firing regimes and cost structures. **F**, No experimental neurons exhibit both high excitability (high *R*_*m*_) and slow adaptation (*τ*_ad_), suggesting functional specialization limits parameter combinations. **G**, Barrel cortex neurons do span the same manifold as visual cortex neurons and not enter the parameter region corresponding to food-restricted animals (red-shaded area, increased *R*_*m*_, depolarized *V*_rest_). **H**, Simulated transitions into the FR regime show a decline in mutual information (top) and energy consumption (bottom) in binary switching tasks (Fig. S11 — see Methods), demonstrating a trade-off where metabolic adaptations save energy at the cost of coding performance.

When under metabolic stress from food restriction, neuronal parameters in the visual cortical dataset alter in a predictable manner. Neurons from energy-deprived animals exhibit an increased membrane resistance and a more depolarized resting potential compared to control animals (Padamsey *et al*., 2022) (Fig. 1A). These changes are consistent with a shift toward energy conservation (Fig. 3), but the neuronal parameters remain on the predicted manifold. The neurons appear to dynamically adjust their parameters depending on energy availability whilst moving along the Pareto front.

**Figure 3.**
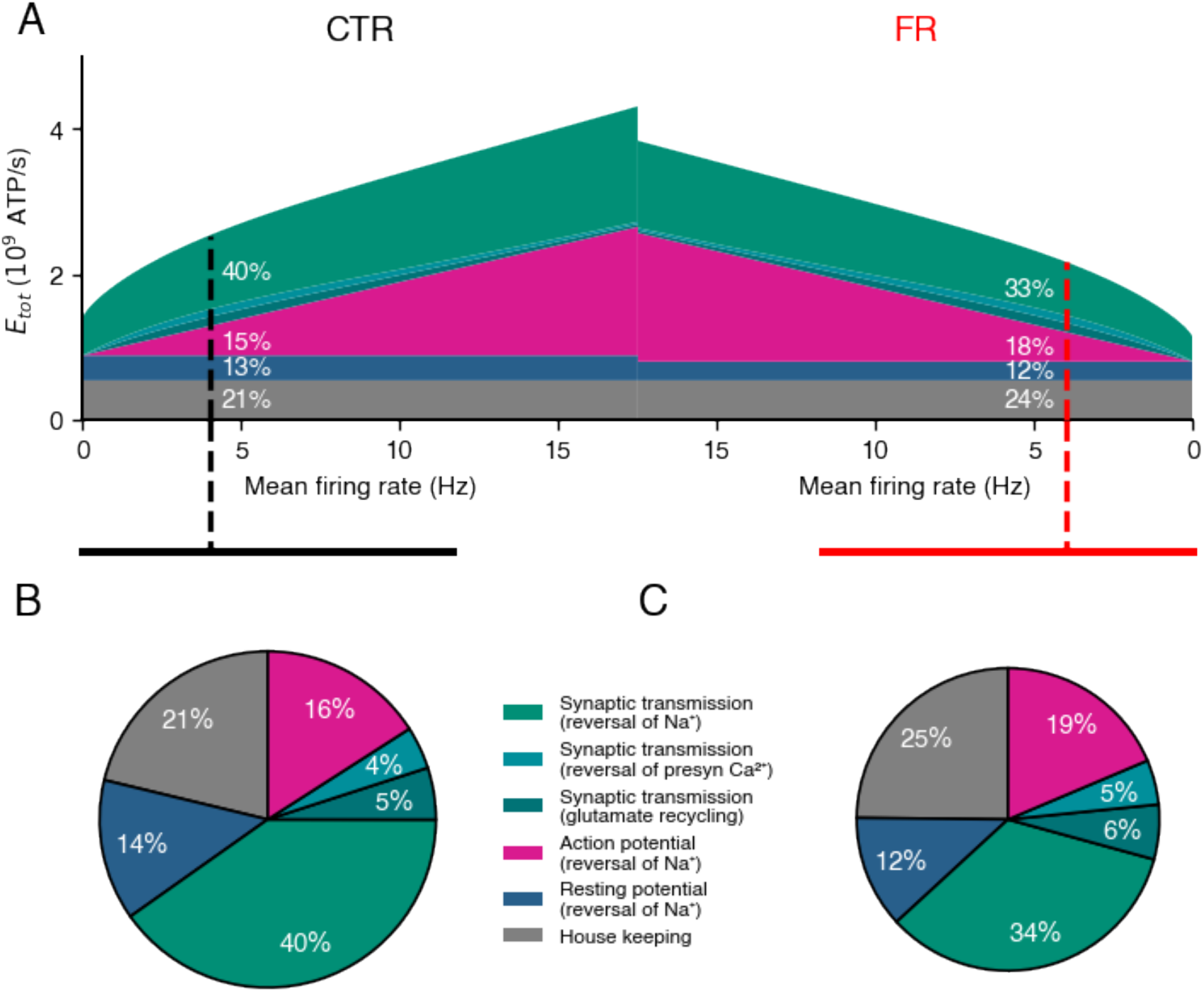
New refined neuronal energy budget estimate with synaptic and non-synaptic contributions and rate-dependence. **A**, Updated energy budget (for derivation see Methods) of cells from control (CTR), left, and food restricted (FR), right, animals with six energy contributors. Cells from food-restricted animals show smaller total energy demand for all firing rates. **B**, CTR and **C**, FR energy components ratios projected out for the “optimal” firing rates of 4 Hz as indicated in the pie charts. They show different ratios of energy components between cells from CTR and FR animals.

### Sensory neurons operate on their Pareto-optimal frontier

Sensory neurons must balance two fundamental objectives: energy efficiency and information transmission. Using a Pareto-optimality framework, we show that neurons from distinct cortical areas populate distinct Pareto fronts that each reflect this fundamental trade-off (Fig. 1B and Fig. 2A,B). In the visual cortex, neurons lie at the Pareto-front spanned by the energy efficiency and information selectivity (OSI) objectives (Fig. 1D), while in the somatosensory cortex, neuronal feature combinations are distributed along a front spanned by energy efficiency and mutual information (MI) (Fig. 2C,D). However the parameters of excitatory and inhibitory cells in somatosensory cortex lie on distinct manifolds. These observations suggest that both area-specific demands (visual vs somatosensory) and cell-type-specific properties (excitatory vs inhibitory) shape distinct energy–information trade-offs.

Depending on functional requirements, sensory neurons may prioritize information accuracy or reduced metabolic cost. For example, neurons processing rapidly changing inputs may optimize for information fidelity, whereas neurons in energetically constrained environments may shift toward more energy-efficient configurations (Remme *et al*., 2018). In line with this, neurons from visual cortex of food-restricted (FR) animals shift away from the high-information region occupied by neurons from control (CTR) animals and toward the energy-efficient regime (Fig. 1D). These energy-saving changes increase the membrane time constant (by an increased membrane resistance), which strongly reduces the ability of the neurons to encode changing stimuli but only slightly impairs orientation selectivity. In contrast, somatosensory neurons — which did not experience metabolic stress — do not populate the energy-conserving parameter space explored by neurons from FR animals (Fig. 2G). Simulations with binary switching tasks (see Methods) reveal that shifting parameters in this direction (as observed by (Padamsey *et al*., 2022): increased membrane resistance, depolarized resting potential, and reduced synaptic drive) does reduce energy demand but simultaneously decreases the information transmission (MI) substantially (Fig. 2H).

Together, these results indicate that, despite a diversity in parameters, neurons lie along an energy–information Pareto frontier. The precise frontiers spanned by different neuronal types in different contexts reflect their functional adaptation to the competing constraints they experience biologically.

### Firing rates are tuned to maximize energy-efficient coding

The coding strategies employed by different neurons also reflect their energetic constraints (Lennie, 2003). Sparse representations reduce the number of spikes required for information transmission, maximizing computational efficiency (Olshausen and Field, 1996) and energy efficiency (Levy and Baxter, 1996; Sengupta et al., 2010). Levy and Baxter (1996) and Laughlin and Sejnowski (2003) made the assumption that neurons maximize their representational capacity per unit of energy expended to predict that firing frequencies should be restricted to optimal ranges of around 6 Hz. Across cortical areas and cell types, we identify a consistent range of optimal firing rates that maximize coding-efficiency (Fig. 1E, Fig. 2E) prioritizing information transmission over energy-efficiency (reducing their *MI/E*_*tot*_ ratio compared to simulation results — Fig. 1F). This range corresponds to the low-to-medium firing rates commonly observed in the mammalian neocortex.

Notably, the exact position and shape of the coding-efficiency curves (Fig. 1E, Fig. 2E) show distinct differences. This suggests that neurons are tuned to area- and cell-type-specific coding demands, reflecting divergent computational roles. Moreover, visual and somatosensory neurons display variations in the steepness and width of their optimal efficiency curves, suggesting different tolerances to firing rate deviations, thus, indicating differences in coding robustness and flexibility. Within barrel cortex, inhibitory neurons appear suited to encode information at higher rates than excitatory neurons (Fig. 2E). Furthermore, barrel cortex neurons do not exhibit combinations of high excitability and slow adaptation — reflected in the absence of cells with both a high membrane resistance (*R*_*m*_) and long adaptation time constants (*τ*_ad_) (Fig. 2F). This constrained distribution likely reflects functional specialization, limiting biophysical variability to support task-specific demands.

We propose that neurons homeostatically regulate their intrinsic excitability and post-synaptic weights to maintain firing rates within their optimal range — consistent with mechanisms of activity-dependent adaptation (Turrigiano et al., 1994; Desai *et al*., 1999; Turrigiano and Nelson, 2004). Padamsey *et al*. (2022) observed that the food-restricted visual cortical neurons did not have significantly different firing rates to the control case. Self-tuning not only maximizes information efficiency but also contributes to population sparsity, a hallmark of efficient coding at the network level.

### Refined neuronal energy budget with rate-dependence allows elevated firing rates in food-restriction being more energy-efficient than controls

To understand neuronal energy expenditure more precisely, we introduce a refined energy budget model (for cortical pyramidal neurons) that builds upon previous estimates (Attwell and Laughlin, 2001; Howarth et al., 2012; Yu *et al*., 2017). Unlike these previous models, which assumed constant firing rates (4 Hz), our approach accounts for rate-dependent energy consumption, adaptation currents, non-spiking regimes, and additional ionic conductances (*I*_*h*_). We demonstrate that including an additional conductance for hyperpolarization-activated cyclic nucleotide–gated (HCN) channels allows a neuron to increase its resting potential while reducing the ATP consumption by an additional ∼ 25% (Fig. S8). Using a detailed multi-compartment neuronal model (Goetz *et al*., 2021), we derive an improved approximation of the energetic cost of non-specific excitatory synaptic currents, enabling us to estimate the instantaneous energy state of integrate-and-fire neuron models (Fig. 3 — see Methods). This refined energy budget reveals that even the modestly (though not statistically significantly) elevated firing rates observed in food-restricted neurons (Padamsey *et al*., 2022) are in fact energetically favorable. Although food-restricted neurons fire slightly more frequently compared to control, their substantially reduced excitatory synaptic currents lead to an overall decrease in total energy consumption. At the same time, the increase in firing rate partially compensates for reduced synaptic drive by preserving information transmission. Consequently, a small upward shift in firing rate represents an efficient trade-off: it maximizes information transfer for only a marginal energetic cost, allowing neurons to operate near an optimal information–energy balance under metabolic constraints.

### Neurons follow predictable trajectories in response to energy deprivation

To understand how neurons might respond dynamically to metabolic stress, we analyzed food-restricted neuronal cell data (Padamsey et al., 2022) and simulated parameter adjustments under energy constraints (see Methods). We assume that neurons during energy shortages try to reduce their energy use, while still maintaining their coding abilities as well as possible. Maintaining the highest possible OSI among the neighbouring parameter combinations that allow lower energy expenditure predicts systematic trajectories in parameter space that:

- reduce excitatory synaptic input currents to conserve energy,
- increase membrane resistance and depolarize the resting potential to enhance excitability or hyperpolarize for small resting energies,
- maintain low firing rates to minimize the energetic cost of action potential generation and neurotransmitter release.

From any combination in the three dimensional parameter space (Fig. 1B) the trajectory moves into the optimal information-per-energy manifold, minimizing the total energy consumption and maintaining a near-optimal energy-information trade-off (Fig. 4). Once in this manifold the trajectories move along this manifold to combinations of high input resistances and depolarized resting potentials maintaining nearly constant firing rates as observed experimentally (Padamsey *et al*., 2022). This confirms that biophysical parameters are strongly constrained by functional needs, while energy is minimized as long as function is maintained (Remme *et al*., 2018).

**Figure 4.**
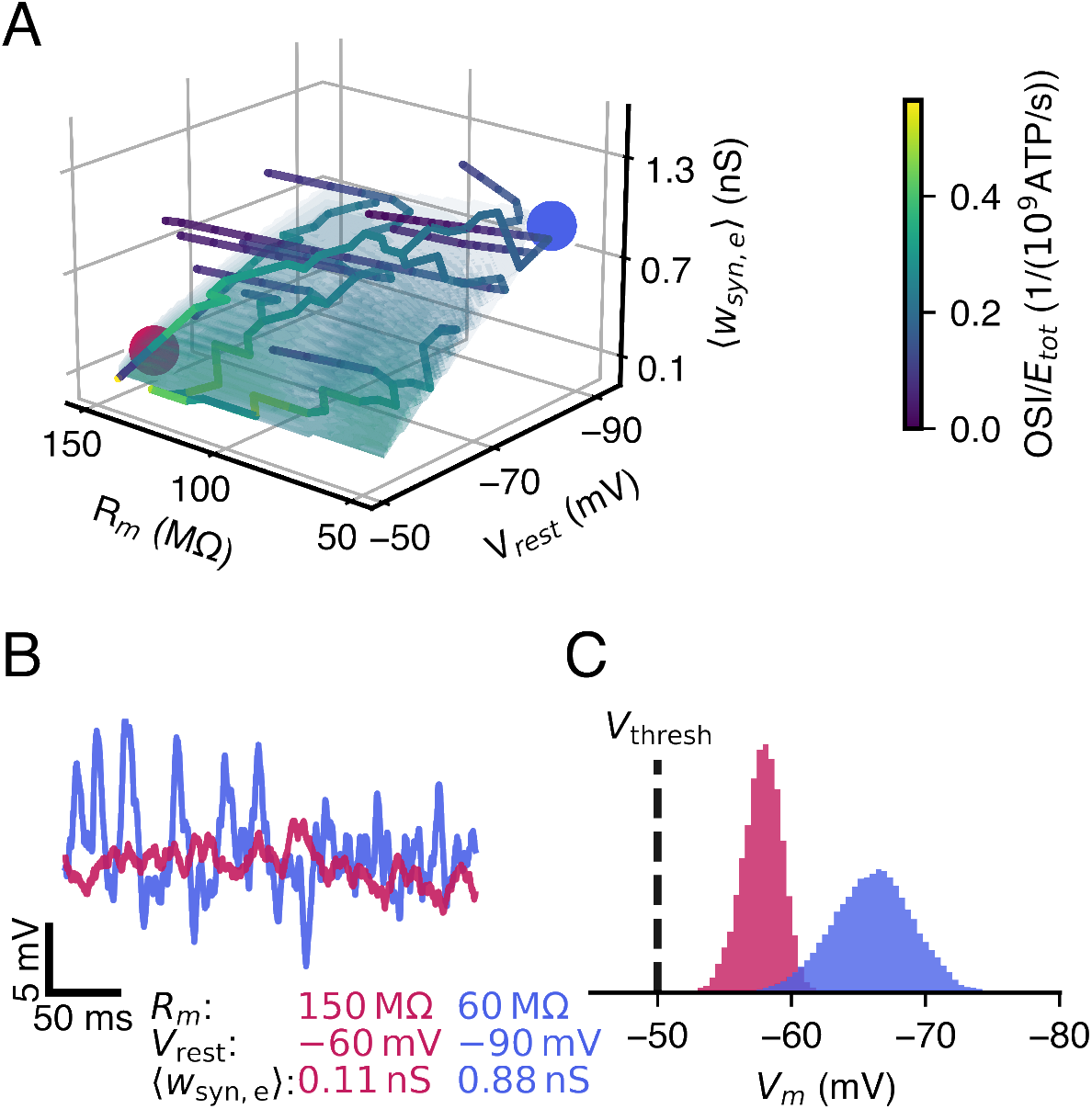
Energy-deprived neurons follow a constrained adaptation trajectory, minimizing their energy expenditure, while maintaining their coding. **A**, Model neurons (Fig. 1B, AdExp model with an orientation selectivity task (Fig. S1) — see Methods) facing energy scarcity adjust their parameters to maintain an optimal energy-information trade-off. Trajectories in the three-dimensional parameter space (membrane resistance, resting potential, total synaptic weight) found altered in food-deprivation experiments (Padamsey *et al*., 2022) first converge onto the predicted high-efficiency manifold, then shift within this manifold towards configurations that reduce energy consumption while preserving information transmission. **B**, Blue and purple traces illustrate example voltage responses before and after adaptation; the cells have preserved spiking activity (0.09 Hz vs 0.11 Hz) despite metabolic constraints. **C**, Normalized histogram of membrane voltages reveals a narrower distribution for depolarized resting potentials, while hyperpolarized resting potentials exhibit broader variability.

The adaptive changes of membrane resistance, resting potential, and synaptic weights align with prior findings that neurons across multiple systems optimize energy efficiency (transmit maximal information per energy used) in axonal transmission (Levy and Baxter, 1996) as well as synaptic transmission (Lezmy et al., 2021; Harris et al., 2012) with presynaptic transmitter release (Levy and Baxter, 2002; Savtchenko et al., 2013; Yuan et al., 2018) and the postsynaptic conductance (EPSCs) (Harris *et al*., 2015, 2019). This supports the notion that energy constraints drive the evolution and development of neuronal biophysics (Niven and Laughlin, 2008), with neurons self-regulating their metabolic expenditure while continuously preserving functional output.

### Non-multiplicative synaptic scaling, membrane noise, and adaptation currents jointly contribute to tuning curve broadening during energy scarcity

We have seen that food restriction results in broader neuronal tuning curves (Padamsey *et al*., 2022) — but there are multiple potential mechanisms that could drive this effect. To investigate its cause, we first inspected the observed synaptic scaling behavior. We examined whether synaptic weight scaling in energy-deprived neurons is multiplicative or non-multiplicative. Fitting the experimental mEPSC distributions (Padamsey et al., 2022) with log-normal functions (Turrigiano *et al*., 1998; Buzsaki and Mizuseki, 2014) revealed non-multiplicative weight scaling (Kolmogorov-Smirnov-test, *D* = 0.44,*p* = 0.003^∗∗^) (Fig. 5A). The non-multiplicative scaling disproportionately weakens strong synapses, reducing overall excitatory drive while preserving firing rates at lower metabolic cost. This is energetically favorable: neurons with non-multiplicative scaling reduce ATP consumption significantly because only a few changes in strong synapses are needed to control the mean firing rate: 1.75 · 10^9^ ATP/s in multiplicative scaling (Fig. 5B) vs. 1.69 · 10^9^ ATP/s in non-multiplicative scaling (Fig. 5C). However, downscaling stronger synapses disproportionately reduces the variance of the synaptic inputs resulting in a reduced dynamic range (Fig. S1E). This also decreases the dynamic range of the membrane potential, which degrades stimulus selectivity, as seen in the broader tuning curves of energy-deprived neurons (Fig. 5B–C) and overall affects the coding abilities. Extrapolating non-multiplicative synaptic weight scaling based on two experimental conditions (CTR and FR), however, comes with high uncertainties. Thus, as a conservative and well-established assumption, we applied multiplicative weight scaling in our other models (Turrigiano *et al*., 1998); in reality, stronger synapses are downscaled disproportionately more than predicted by a strictly multiplicative model. Consequently, we expect the actual effects to be more pronounced.

**Figure 5.**
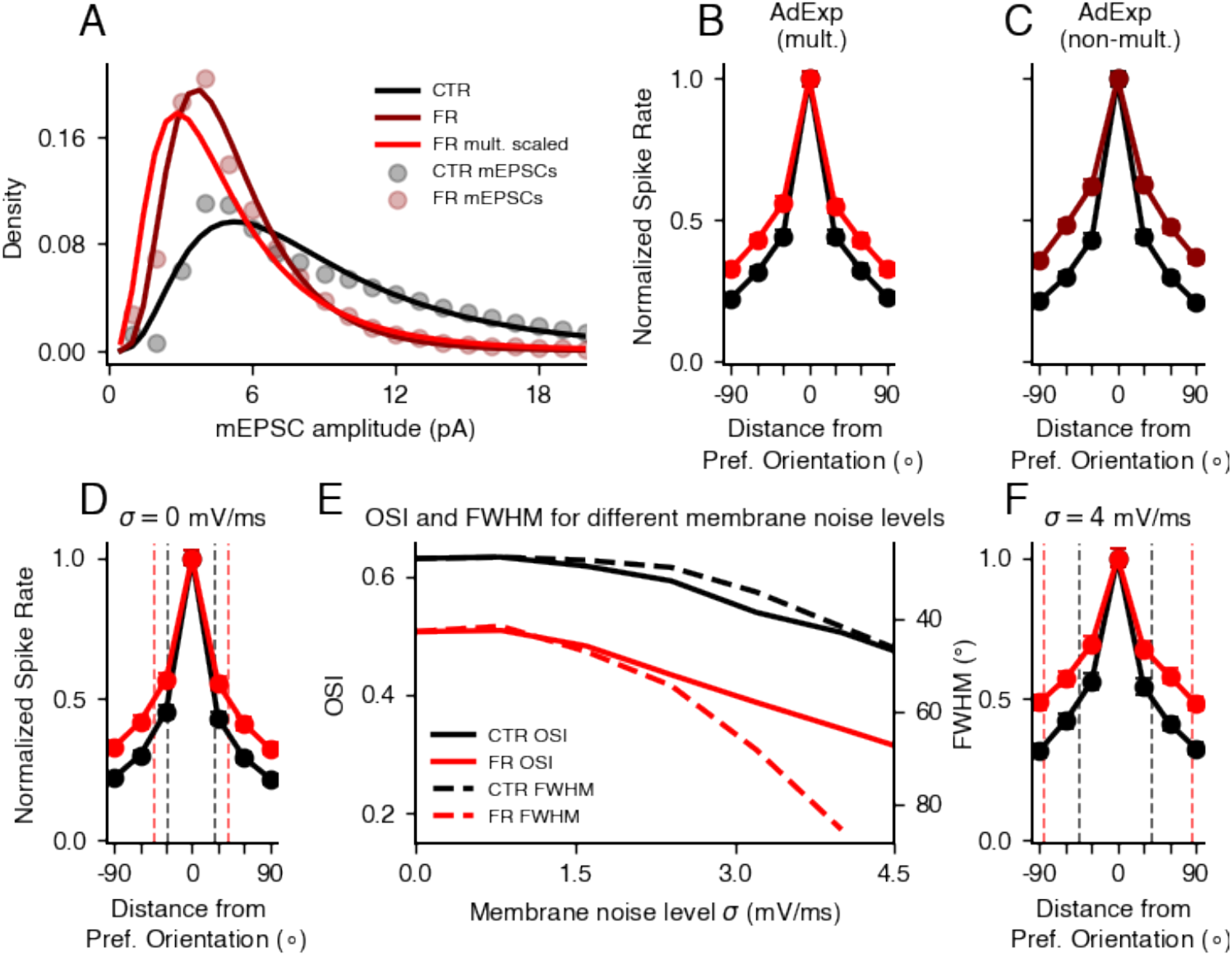
Tuning curve broadening under energy deprivation is amplified by non-multiplicative synaptic weight scaling and membrane noise. **A**, Experimental *ex vivo* mEPSC distributions of neurons from control (CTR, black) and food-restricted (FR, dark red) animals (Padamsey *et al*., 2022). A multiplicative downscaling of CTR weights (red) (Turrigiano et al., 1998) fails to reproduce the experimental FR distribution, indicating a non-multiplicative scaling mechanism (dark red). CTR neurons show a distribution with mean *µ*_CTR_ = 9.70 and standard deviation *σ*_CTR_ = 6.92, while FR neurons have a lower mean *µ*_FR_ = 5.26 and narrower spread *σ*_FR_ = 2.75. Multiplicatively scaling the CTR distribution to match the FR mean yields *µ*_CTR,scaled_ = 5.26 and *σ*_CTR,scaled_ = 3.75, indicating that the FR distribution cannot be explained by multiplicative scaling of the CTR distribution. **B**, Simulated tuning curves (AdExp model with an orientation selectivity task (Fig. S1) — see Methods) using multiplicative weight scaling (red) show broadening. **C**, In contrast, non-multiplicative downscaling of synaptic weights (dark red) shows more pronounced broadening, suggesting that strong synapses are preferentially weakened — reducing selectivity while improving energy efficiency. **D**, Simulated tuning curves without intrinsic membrane noise show smaller broadening compared to high membrane noise levels as in **F**. CTR neurons retain higher orientation selectivity (OSI, solid lines) and narrower tuning (lower full width at half maximum (FWHM), dashed lines) than FR neurons. **E**, Orientation selectivity index (OSI, solid) and full width at half maximum (FWHM, dashed) as a function of membrane noise amplitude. Solid lines show a monotonic decline in OSI for CTR as well as FR but with consistently lower values in the FR case. Dashed lines show an increased decline in FWHM for FR compared to CTR representing a loss in selectivity of FR neurons. Errorbars are within width of line. **F**, Tuning curve at high membrane noise (4 mV/ms) illustrates strong broadening and loss of selectivity in FR neurons, with markedly increased FWHM (dashed lines) compared to CTR neurons.

A second factor contributing to tuning curve broadening is intrinsic membrane noise, with neurons from food-restricted (FR) animals consistently showing lower orientation selectivity than control (CTR) across all tested noise levels (Fig. 5E) when presented different oriented gratings (Fig. S1A–D). This reduction in selectivity is reflected in broader tuning curves (Fig. 5D–F) and becomes more pronounced as noise increases. The resulting heightened sensitivity of FR neurons to intrinsic noise likely stems from their reduced voltage gap between resting and threshold potentials, making them more susceptible to random fluctuations. In addition, experimental *ex vivo* data show rather constant levels of membrane noise in FR neurons (0.9 mV/ms) and controls (0.8 mV/ms) (Fig. S9A-C).

Finally, we find that adaptation currents play a crucial role in tuning curve broadening. Simulations with an oriented grating inputs (Fig. S1 — see Methods) using a leaky integrate-and-fire (LIF) model show minimal broadening, whereas the adaptive exponential integrate-and-fire (AdExp) (Brette and Gerstner, 2005; Zerlaut and Destexhe, 2017) model captures the experimentally observed effects (Fig. S3). We propose that subthreshold and spike-triggered adaptation currents suppress strong responses to preferred stimuli while maintaining overall activity levels. Overall, this reduces orientation selectivity.

### Resting potential adaptation enhances coding efficiency and is optimized *in vivo*

Why do neurons adjust their resting membrane potential rather than their spiking threshold in response to metabolic stress, such as during food restriction (Padamsey *et al*., 2022)? To address this, we compared scenarios in which the voltage gap between resting and threshold potential was modulated either by depolarizing the resting potential or by lowering the spiking threshold (AdExp model with an orientation selectivity task (Fig. S1) — see Methods). These two options are not symmetrical, as both excitatory and inhibitory synaptic reversal potentials remain fixed. Altering the range of voltages a neuron can experience below threshold will therefore change the strength and integration of synaptic inputs. Across a range of gap values the orientation selectivity index (OSI) was consistently higher when the resting potential was adapted (Fig. 6A – Top). By contrast, threshold adaptation led to a rapid decline in OSI and an increase in firing rate (Fig. S10A), reducing coding efficiency (Fig. S10E) and increasing metabolic cost (Fig. 6A – Bottom). These results indicate that shifting the resting potential maintains selectivity at lower energetic expense.

**Figure 6.**
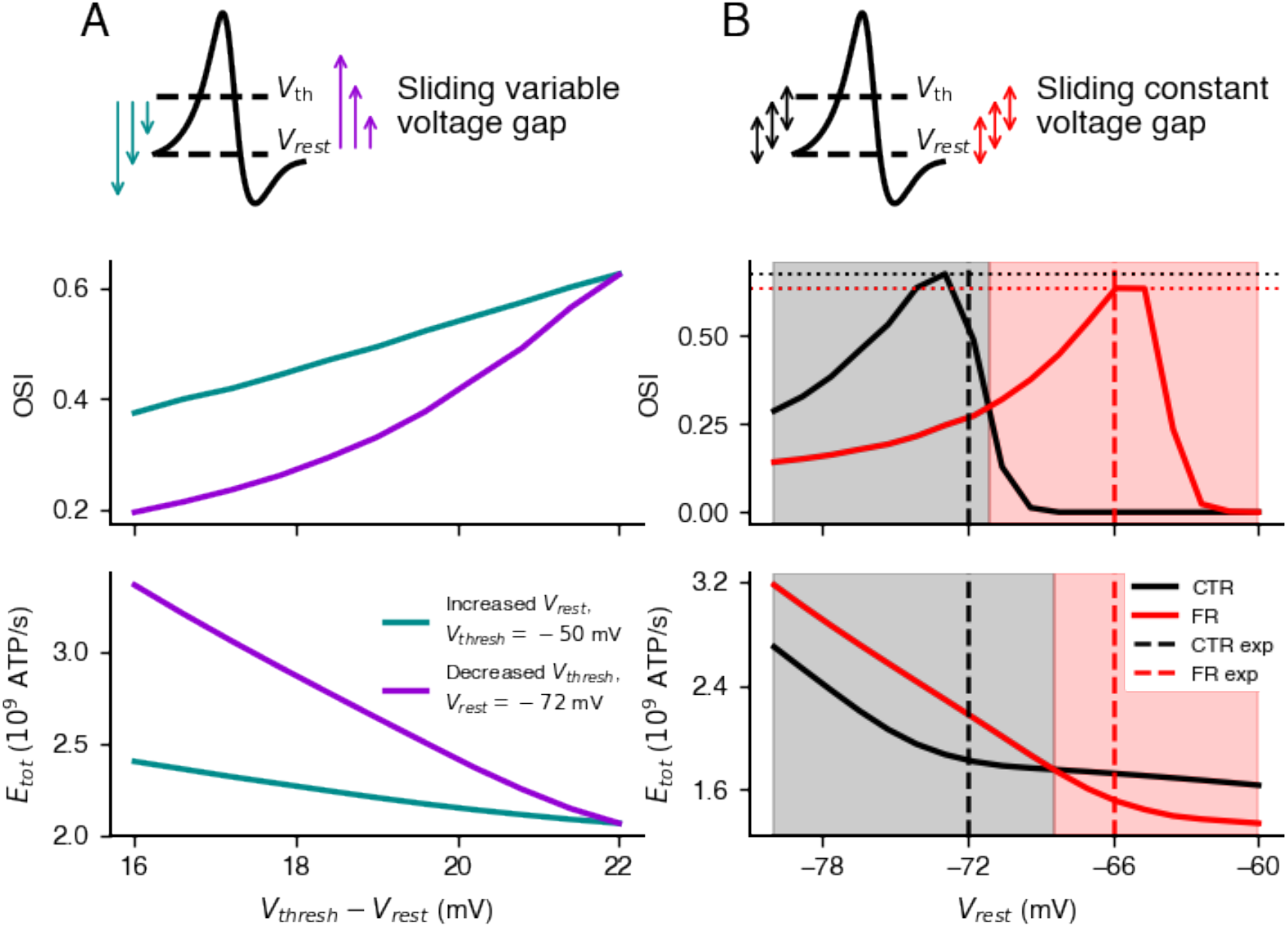
Resting potential adaptation enhances coding performance and energy efficiency. **A**, Comparison between two mechanisms of increasing excitability — raising the resting potential (turquoise) versus lowering the threshold (purple) — for a range of voltage gaps in orientation selectivity tasks (Fig. S1 — see Methods). Errorbars are within width of line. **A – Top**, Orientation selectivity index (OSI) is consistently higher when excitability is increased via resting potential depolarization. **A – Bottom**, Total energy expenditure (*E*_*tot*_) is markedly higher when reducing the threshold, driven by elevated firing rates (Fig. S10A). **B**, Coding performance and energy cost as a function of resting membrane potential, while keeping the rest–threshold voltage gap constant (35% synaptic weight reduction in FR relative to CTR, based on experimental data, using an experimentally measured membrane noise level of 0.9 mV/ms). CTR neurons have a 6 mV wider gap than FR neurons. **B – Top**, Orientation selectivity index (OSI) across a range of resting potentials. Coding performance peaks near the experimentally observed resting potentials for cells of control (–72 mV and food-restricted (–66 mV) animals (dashed lines) (Padamsey et al., 2022). CTR neurons achieve higher overall selectivity. Shaded areas indicate optimal coding regimes for CTR (black) and FR (red) neurons. **B – Bottom**, Total energy consumption over the same resting potential range. Energy efficiency closely follows coding performance: CTR and FR neurons are most efficient near their respective *in vivo* resting potentials.

To further investigate this mechanism, we systematically varied the resting potential while maintaining a fixed voltage gap to threshold, spanning the range between inhibitory and excitatory synaptic reversal potentials. This revealed a distinct coding optimum — reflected in the orientation selectivity index (OSI) — for specific resting potentials (Fig. 6B – Top) with a higher overall selectivity in the control case. Strikingly, these optima coincide with the experimentally observed resting potentials in control and food-restricted animals (CTR: –72 mV, FR: –66 mV; dashed lines in Fig. 6B). Moreover, energy consumption tracked coding performance: control neurons were most efficient near their mean experimental resting potential, and food-restricted neurons around their mean resting potential (Fig. 6B – Bottom). Notably, transitioning from CTR to FR parameters shifts neurons toward lower overall coding capacity (indicated by dotted lines), but yields an improved energy efficiency. These findings suggest that cortical neurons dynamically tune their resting membrane potential to operate within a biophysically optimal regime that is coding-efficient as well as metabolically efficient.

Experimental data (Padamsey *et al*., 2022) show variability in spiking thresholds, but mean threshold values remain unchanged under food restriction. Instead, neurons adjust their resting potentials to optimize energy efficiency. This suggests that resting potential modulation is a primary mechanism for energy adaptation, while threshold adjustments may serve different functional roles. In previous work we have found that sub-threshold adaptation of the *h*-current increases information transfer when inputs vary in time (Zeldenrust et al., 2024). Depression of the *h*-current leads to more false alarms and decreases information transfer rates. On the other hand, threshold adaptation increases the working range of neurons. This was not tested for in the food-restriction experiments (Padamsey *et al*., 2022), but might represent another important difference in how neurons adapt to optimally represent the stimuli they receive *in vivo*.

## Discussion

We have demonstrated that there is hidden structure in the diversity of neuronal properties reflecting an energy-information trade-off. We have developed an energy budget model that accounts for neuronal energy expenditure based on firing rate and intrinsic neuronal properties (see Methods). Using this model we have found that neurons control their intrinsic properties to populate a neuron type- and context-dependent Pareto-optimal space, balancing energy efficiency and information transmission. This trade-off predicts a low to medium firing rate regime as observed in neocortex as particularly efficient. It also predicts systematic adaptations under energy scarcity, favouring more energy-efficient configurations at the expense of a relatively small information loss. While we have revealed a hidden structure in the space of neuronal properties, reflecting neuronal degeneracy, it is important to ask why there is so much diversity in neuronal parameters in the first place.

Heterogeneity of intrinsic neuronal properties is ubiquitous in the brain and cells of the same cell type show diverse structures and response properties (Altschuler and Wu, 2010; Joshi *et al*., 2025). It is conceivable that this diversity simply reflects randomness during neuronal development. However, computational work has shown that such heterogeneous populations of neurons improve the performance of neural networks by increasing the variability in the representation of stimulus features (Duarte and Morrison, 2019; Shen et *al*., 2023) resulting in optimized information transmission (Haggard and Chacron, 2023), higher efficiency, and more robust learning relative to networks with homogenous units (Zeldenrust *et al*., 2021; Dahmen et al., 2025). Furthermore, heterogeneity can serve as an additional degree of freedom to reduce time and energy spent on hyperparameter tuning when training artificial neural networks, which is one of the largest costs in practical engineering applications (Perez-Nieves *et al*., 2021).

Biological evidence for benefits of neuronal diversity comes from a study showing that the loss of heterogeneity (difference of resting membrane potential to spiking threshold) is linked to synchronization and epileptic behavior (Rich et al., 2022), highlighting the functional importance of intrinsic variability in biological brains. Heterogeneity of neuronal properties can also be spatially organized. Along the dorsal–to-ventral axis of the hippocampus, CA1 pyramidal neurons (Dougherty *et al*., 2012) and dentate granule cells (Kumari and Narayanan, 2026) (with similar gradients reported in medial entorhinal cortex stellate cells) exhibit a systematic increase in input resistance and a concomitant depolarization of resting membrane potential. It is currently unclear what the functional benefit of this may be. Based on our findings we propose that these gradients reflect energy-efficient solutions to gradually changing information processing needs along this axis.

Energy efficiency has long been identified as a principle of nervous system organization (Barlow, 1961). This principle has been applied at various levels ranging from synaptic transmission (Goldman, 2004; Harris et al., 2012) to information coding at the population level Barlow (1961). In this study, we have focused on the question of how coordinated changes in neuronal parameters affect information coding in single neurons response to sensory stimuli. Our work complements earlier studies showing that balanced excitatory and inhibitory currents give rise to more energy-efficient (bits/ATP) and informative (bits/spike) coding (Sengupta et al., 2013; Yu et al., 2018).

Biological brains have to deal with diverse changes in energy supply. Food scarcity induces systematic changes in resting potential and membrane resistance, affecting neuronal integration properties (Padamsey et al., 2022). An increased membrane time constant alters information integration. This might have big effects in tasks such as coincidence detection where high temporal precision is needed (Remme *et al*., 2018). Changes in the distribution of time constants could impact network-level task performance, its learning and robustness (Perez-Nieves *et al*., 2021; Shen et al., 2023).

While such adaptations to food scarcity take weeks, sensory cortices may be able to modify their energy-information trade-off on much faster time sacles. Recent findings suggest that inhibitory circuits dynamically regulate coding precision and perception (Gauld *et al*., 2024). The authors show that circuit properties and cell intrinsic properties work together to form a “latent” pool of neurons that is normally silenced by sensory-evoked inhibition, but that can be artificially recruited via optogenetic stimulation to enhance perception. When activated, these neurons increase the encoding quality of a stimulus. These findings suggest that sensory cortices may be able to rapidly control their energy-information trade-off by altering the level of inhibition.

Interest in the energetic supply and demand of neurons has grown substantially in recent years due to the recognition that metabolic dysfunction is tightly linked to neurological disease (Bazzigaluppi *et al*., 2017), e.g., in the case of mitochondrial dysfunctions (Klemmensen *et al*., 2024). Although neurons are astonishingly flexible when handling transient energetic mismatches, chronic metabolic perturbations have detrimental effects on neuronal function, ultimately contributing to the development of neurodegeneration (Köhler-Solís and Schirmeier, 2025). Potentially, a long-term energy depletion results in a chronic depolarization and drifting reversal potentials making a network prone to seizures (Le Masson *et al*., 2014). Our framework could provide insights into how neurons adapt to chronic energy deficiencies, such as those observed in Alzheimer’s disease (Engl and Attwell, 2015).

Evolution has resulted in many ways of improving the energy efficiency of nervous systems. An improved understanding of the various mechanisms can also inspire engineering efforts to build more energy-efficient artificial computing systems. This has long been a major motivation behind the field of neuromorphic engineering (Indiveri, 2025). For example, recent approaches have considered mechanisms for reducing the energy consumption of encoding information (Zhang et al., 2025; Dimakou *et al*., 2025; N’dri et al., 2025) or learning (Li and van Rossum, 2020; Pache and van Rossum, 2023). We believe that engineers still have much to learn from how biological nervous systems flexibly manage their energy budgets in the face of fluctuating supply and demand.

## Conclusion

By integrating experimental data with computational modeling, we have identified degenerate solutions to the problem of neuronal parameter selection revealing an optimal energy-information trade-off. Our findings suggest that neurons operate within a constrained subspace of parameter combinations that ensure close to optimal information transmission. Our model predicts a high but well-orchestrated variability of neuronal parameters (reflecting degeneracy), which is context and cell-type-specific. Under energy limitation, neurons systematically shift their parameters along a Pareto-optimal frontier, reducing metabolic costs while preserving functional output as much as possible. These results provide a theoretical framework for understanding neuronal degeneracy, metabolic adaptation, and functional diversity across cortical areas. Our work highlights energy efficiency as a fundamental principle of neural computation, offering new insights into how neurons balance metabolic demands with information processing needs. Overall, efficient energy management presents itself as a promising research area spanning biological adaptation, disease mechanisms and technological design.

## Acknowledgments

We want to thank Zahid Padamsey (Weill Cornell Medicine, Qatar) for his provision and explanation of the experimental data, as well as his feedback on our results. We are grateful to Arnd Roth (University College London) for his advice and expertise on detailed compartmental models and energy estimation. We are thankful to Richard Gao (Goethe University Frankfurt) for his advice on the SBI pipeline.

## Author contributions

P.S., A.D.B., P.J., J.T. designed the study.

P.S. and A.D.B. performed the simulations and analyzed the data.

P.S., F.Z., A.D.B., P.J., J.T. wrote the paper.

F.Z. supervised the experiments.

We reanalyzed the experimental data from a previous study (Zeldenrust et al., 2017). In this study Fleur Zeldenrust and Tansu Celikel supervised the experiments. Fleur Zeldenrust designed the experimental protocol. Fleur Zeldenrust and Niccolo Calcini did the experimental data analysis. Niccolo Calcini, Koen Kole, Ate Bijlsma, Xuan Yang performed the experiments.

During the preparation of this work the authors used ChatGPT-5 for inspirational language work, improvements to human-generated texts for readability and style, and to ensure that the texts are free of errors in grammar, spelling, punctuation, and tone as well as the readability and debugging of code. After using this tool, the authors reviewed and edited the content as needed and take full responsibility for the content of the published article.

## Funding

P.S. was supported by Studienstiftung des Deutschen Volkes.

F.Z. acknowledges funding from NWO (a Vidi grant VI.Vidi.213.137 and “DBI2” (024.005.022), a Gravitation program of the Dutch Ministry of Science, Education and Culture).

J.T. was supported by the Deutsche Forschungsgemeinschaft (DFG, German Research Foundation) through the Excellence Cluster EXC3066 “The Adaptive Mind” and the priority program SPP 2041, project numbers TR 881/7-2; TR 881/8-2, priority program SPP 2411, project numbers 520617944; TR 881/11-1 and Research Unit FOR 5368, DYNAMIC center, funded by the LOEWE program of the HMWK (grant number: LOEWE1/16/519/03/09.001(0009)/98

J.T. acknowledges support from the Johanna Quandt foundation.

The funders had no role in the study design, data collection, analysis, or decision to publish.

### Declaration of interests

The authors declare no competing interests.

## Material and methods

### Experimental data

The raw experimental data is accessible by the original publications (Padamsey *et al*., 2022; Zeldenrust et al., 2024). Extracted data and example data is available with the analysis code on github (https://github.com/trieschlab/energy-information-trade-off).

#### Energy-deprivation data from Padamsey *et al*. (2022)

Padamsey *et al*. (2022) investigated the effects on food deprivation on layer 2/3 pyramidal neurons in visual cortex.

##### *In vivo* response to oriented gratings

Padamsey *et al*. (2022) presented drifting gratings (12 different angles from 0° to 330°, 30° increments) with a temporal frequency of 1 Hz for 1.5 s in random order followed by a 1 s grey screen. The authors measured following neuronal parameters via *in vivo* electrophysiology:

- Current clamp recordings: 33 control and 26 food-restricted animals
- Voltage clamp recordings: 13 control and 10 food-restricted animals.

The key findings include:

- an 18 % mean increase in membrane resistance *R*_*m*_,
- a 10 % mean increase in resting membrane potential *V*_rest_,
- a 42 % mean reduction of synaptic currents (excitatory and inhibitory),
- a constant spiking threshold *V*_th_,
- a constant spiking rate *r*_post_,
- a constant E-I ratio as quantified by the synaptic currents.

The *in vivo* electrophysiological measurements of a single cell contains its membrane resistance *R*_*m*_, its resting membrane potential *V*_rest_ and its spiking rate *r*_post_. For comparison with our simulations, the synaptic scaling parameter *w*_scale_ was estimated for each experimental condition by fitting experimental firing rates to simulated firing rates integrating 5000 homogeneous Poisson inputs. The resulting synaptic weights were then scaled by a factor to match experimental and simulated tuning curves of the non-homogeneous firing of oriented grating presentation.

##### *Ex vivo* measurements of synaptic weight distributions

In *ex vivo* electrophysiology measurements of visual cortex neurons (Padamsey *et al*., 2022) recorded AMPAR-mediated mean miniature excitatory postsynaptic currents (mEPSC) at -70 mV in 20 control and 20 food-restricted cells in the presence of 1 mM TTX (Padamsey et *al*., 2022). Results showed:

- a decreased mean mEPSC amplitude,
- a constant mEPSC frequency.

The mEPSC distributions were fitted with log-normal functions as proxies for synaptic weights (Buzsaki and Mizuseki, 2014) (see Fig. 5).

We assume that information stored in synaptic weights can be identified by the average excitatory postsynaptic currents.

#### Estimation of membrane noise

Fluctuations in membrane voltage were estimated from existing *in vivo* blind patch clamp recordings in control and food restricted mice (Padamsey *et al*., 2022). It was first necessary to remove experimental artefacts from the raw traces (Fig. S9A). These artefacts came in three forms: 1) Sudden sharp deviations in voltage. 2) Low-frequency oscillations. 3) Slow baseline drift. 1) Sharp deviations were removed by cutting out 0.2 ms either side of voltages outside the windows [−70 mV, −50 mV] (CTR) and [−90 mV, −65 mV] (FR). 2) Slow oscillations were removed with Notch filters (SciPy Signal Processing package) centered around the first spectral peaks in each case (quality 0.1) (Fig. S9B). 3) Baseline drift was removed by subtracting the moving average (over 0.5 ms) from the voltage. Removing these artefacts gave cleaner estimates of Gaussian membrane voltage noise (Fig. S9C).

#### Simulation-based inference of neuronal parameters of switching task data of Zeldenrust *et al*. (2024)

Zeldenrust *et al*. (2024) measured the intrinsic information encoding properties of excitatory and inhibitory neurons in L2/3 of the mouse barrel (somatosensory) cortex. We found:

- excitatory neurons: high thresholds and strong adaptation resulting in sparse firing and a strong compression of information.
- inhibitory neurons: firing at high rates, resulting in a higher transfer of information.

To infer neuronal parameters from experimental voltage traces and injected currents from Zeldenrust *et al*. (2024), we employed simulation-based inference (SBI) using the `sbi` Python package (Tejero-Cantero et al., 2020). Parameter inference was performed using a two-stage sequential neural posterior estimation (SNPE) framework (Greenberg *et al*., 2019; Lueckmann et al., 2021).

##### Pre-processing and fixed parameters

The resting potential (*V*_rest_) was estimated as the y-intercept of a linear fit between the membrane voltage and the injected current trace. The spiking threshold (*V*_th_) was taken directly from the experimental estimates reported in Zeldenrust et al. (2024). These parameters were fixed throughout the inference procedure.

##### Stage 1: Passive and spike-generation parameters

In the first stage, we inferred parameters governing passive membrane properties and spike initiation: membrane resistance (*R*_*m*_), membrane capacitance (*C*_*m*_), and the spike slope factor (Δ_*T*_). Summary statistics included the mean and standard deviation of the voltage trace, selected voltage percentiles (5, 10, 25, 50, 75, 95%), spike count, and spike-triggered averages.

Priors were defined as:

- *R*_*m*_ ∈ [25, 500] M”
- *C*_*m*_ ∈ [2, 120] pF
- Δ_*T*_ ∈ [0.5, 5] mV

Posterior distributions were estimated using SNPE with direct sampling. If direct sampling exceeded a predefined runtime threshold, posterior sampling was performed using Markov chain Monte Carlo (MCMC) with slice sampling.

##### Stage 2: Adaptation-related parameters

In the second stage, we inferred parameters governing post-spike and adaptive dynamics: reset potential (*V*_reset_), adaptation time constant (*τ*_ad_), subthreshold adaptation strength (*a*_ad_), and spike-triggered adaptation increment (*b*_ad_). Summary statistics included inter-spike interval (ISI) features (mean, standard deviation, coefficient of variation), spike count, first and last spike times, and after-spike averages.

Priors were defined as:

- *V*_reset_ ∈ [*V*_rest_ − 10, *V*_th_ − 1] mV
- *τ*_ad_ ∈ [5, 500] ms
- *a*_ad_ ∈ [1, 10] nS
- *b*_ad_ ∈ [0.001, 0.1] nA

##### Posterior validation and dimensionality reduction

Posterior mean parameter estimates were used to simulate voltage traces. Model quality was assessed by comparing experimental and simulated spike trains using the Earth Mover’s Distance (EMD) (Sihn and Kim, 2019), with data retained for further analysis only if EMD was below 270 ms.

One- and two-dimensional posterior marginals are shown in Fig. S12. Correlations between inferred parameters across neurons are shown in Fig. S13. The quality-checked parameter sets were projected into principal component space (Fig. S14) and used as basis for further simulations.

### Neuron model

We combine simulations of a generic leaky-integrate-and-fire (LIF) neuron model and an adaptive exponential integrate-and-fire (AdExp) neuron model in conjunction with our updated version of the energy calculations based on Attwell and Laughlin (2001); Howarth *et al*. (2012); Yu *et al*. (2017) and information-theoretic measures.

Our simulations employ the leaky-integrate-and-fire (LIF) neuron model (Brunel and van Rossum, 2007), which is a simplified neuron model capturing the essential dynamics of neuronal behavior combined with computational simplicity:

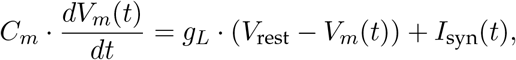

where *C*_*m*_ is the membrane capacitance, *V*_*m*_(*t*) the membrane potential, *g*_*L*_ the leak conductance, *V*_rest_ the leak reversal potential, and *I*_syn_(*t*) the synaptic input current which is modeled by conductance-based excitatory (AMPA) and inhibitory (GABA A) synapses. The neuron spikes if the membrane voltage exceeds the threshold voltage *V*_th_ (*V*_*m*_ *> V*_th_) and is reset to *V*_reset_ after a spike was fired (for some of the simulations we kept it constant at -60 mV but also set it to *V*_rest_ for others to check consistency — see Fig. S4). No qualitative nor significant quantitative changes we observed except for the coefficient variation — see Fig. S4.). The membrane time constant *τ*_*m*_ determines the temporal charge and discharge properties of the membrane and is defined as:

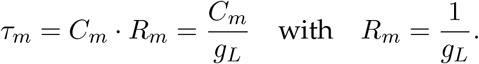

To model biological spiking behavior more realistically the LIF can be extended by an adaptation current and an exponential term for spike initiation, which captures both subthreshold adaptation and spike-triggered adaptation. The adaptive exponential integrate-and-fire (AdExp) neuron model (Brette and Gerstner, 2005) can be represented by the following set of differential equations with the membrane potential dynamics:

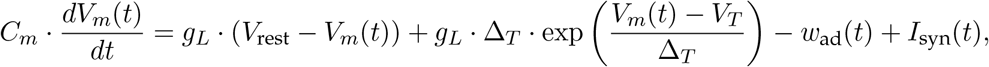

and the adaptation current dynamics:

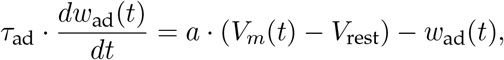

where Δ_*T*_ is the slope factor for the exponential term, *V*_*T*_ the exponential onset voltage, *w*_ad_(*t*) the adaptation current, *τ*_ad_ the adaptation time constant, *a*_ad_ the subthreshold adaptation, and *b*_ad_ the spike-triggered adaptation for which *w*_ad_(*t*) is incremented for each time *V*_*m*_(*t*) reaching the spiking threshold. These differential equations are numerically integrated with built-in functions of Brian2 based on Fourth-order Runge-Kutta algorithms.

#### Stochastic membrane noise

To simulate intrinsic membrane fluctuations, we added a noise term *I*_noise_ to the membrane equation. This current was modeled as uncorrelated Gaussian white noise with zero mean and a standard deviation *σ*_noise_ derived from the experimental *ex vivo* mEPSC recordings. The noise was scaled by the square root of the integration timestep Δ*t* to ensure correct physical units and variance:fd

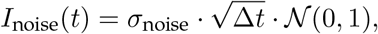

where 𝒩 (0, 1) denotes a standard normal random variable. Due to numerical stability constraints in the presence of stochastic input, simulations including *I*_noise_ were performed using the forward Euler integration scheme instead of the default Runge–Kutta solver in Brian2.

#### Model oriented gratings as synaptic input

The synaptic input current is the sum of all excitatory and all inhibitory synaptic inputs:

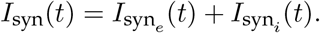

The synaptic inputs are modeled by conductance-based excitatory and inhibitory synapses:

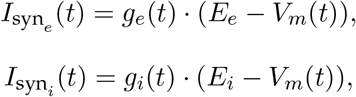

where *g*_*e*_(*t*) is the excitatory synaptic conductance, *E*_*e*_ the reversal potential of excitatory synapses, *g*_*i*_(*t*) the inhibitory synaptic conductance, and *E*_*i*_ the reversal potential of inhibitory synapses. The dynamics of the conductances is described by:

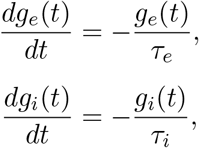

where *τ*_*e*_ is the postsynaptic potential (PSP) time constant for excitatory synapses and *τ*_*i*_ for inhibitory synapses. For every presynaptic spike the respective conductance is increased by the size of the synaptic weight *w*_*e*_ or *w*_*i*_. The excitatory synaptic weights are approximated as log-normally distributed with the *ex vivo* measured mEPSC distribution of the control (CTR) animals with 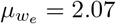 and 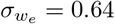 (Padamsey et al., 2022), where the mESPCs are assumed as a proxy for the synaptic weights (Song *et al*., 2005). The same weight distribution is assumed for inhibitory synaptic weights but their total weight is scaled to 25 % of the total excitatory synaptic weight to result in an overall balance of excitation Deneve and Machens, 2016) and inhibition (. The synaptic inputs are excitation-inhibition balanced as experimentally measured (Padamsey *et al*., 2022; Haider *et al*., 2013; Barral and Reyes, 2016). Thus, we maintain in our simulations a constant ratio between excitation and inhibition with the conductance ratio of *g*_*e*_*/g*_*i*_ ≈ 0.52 and the synaptic current ratio of 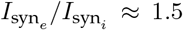 consistent with experimental measurement. The constancy of the excitation-inhibition balance is achieved by scaling excitatory and inhibitory synapses by the same factor *w*_scale_ proportionately (coupling them). For *w*_scale_ = 100 the mean excitatory synaptic weight is *w*_*e*_ = 0.52 nS and the mean excitatory synaptic weight is *w*_*i*_ = 0.4 nS. The excitatory presynaptic firing rates arriving at the synapses are normally distributed with 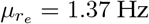 and 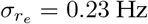 (Goetz *et al*., 2021). The inhibitory presynaptic firing rates are assumed to be uniform with a firing rate of 6 Hz. We included 5000 synapses in total from which 75 % (3750) are excitatory and 25% (1250) are inhibitory (the latter being driven by a homogeneous Poisson process). Of the excitatory synapses 50 % code for a presented stimulus (driven by an inhomogeneous Poisson process) and 50 % are assumed noise synapses (driven by a homogeneous Poisson process) (see Fig. S1C, Fig. S2).

To model *in vivo* like synaptic input of oriented gratings for layer 2/3 pyramidal neurons we assume that stimuli are solely coded in firing rate (rate-based code) and match the firing rates and synaptic weights of the coding synapses. This is motivated by the findings that the connection between neurons with similar receptive fields are stronger (Goetz *et al*., 2021) compared to unrelated receptive field properties. The matching logic is performed similarly to Goetz et al. (2021) (see Fig. S1A–C):

- For the preferred orientation, large synapses are assigned to high firing rates, small synapses with low firing rates.
- For the non-preferred orientation, large synapses fire with low firing rates, small synapses with high firing rates.
- For the intermediate orientations, we interpolate between preferred and non-preferred and draw the firing rates from two Gaussian distributions with same mean and variance.

For simulation purposes different oriented gratings are presented for 1.5 s with 1 s intermittent background firing and with the same increments as in the food-deprivation experiments (Padamsey *et al*., 2022) following the scheme 0°, 30°, 60°, 90°, 120°, 150°, 180° with 90° as the preferred stimulus and 0° and 180° as the non-preferred stimulus (Fig. S1A). For more references check Goetz *et al*. (2021).

**Table 1:**
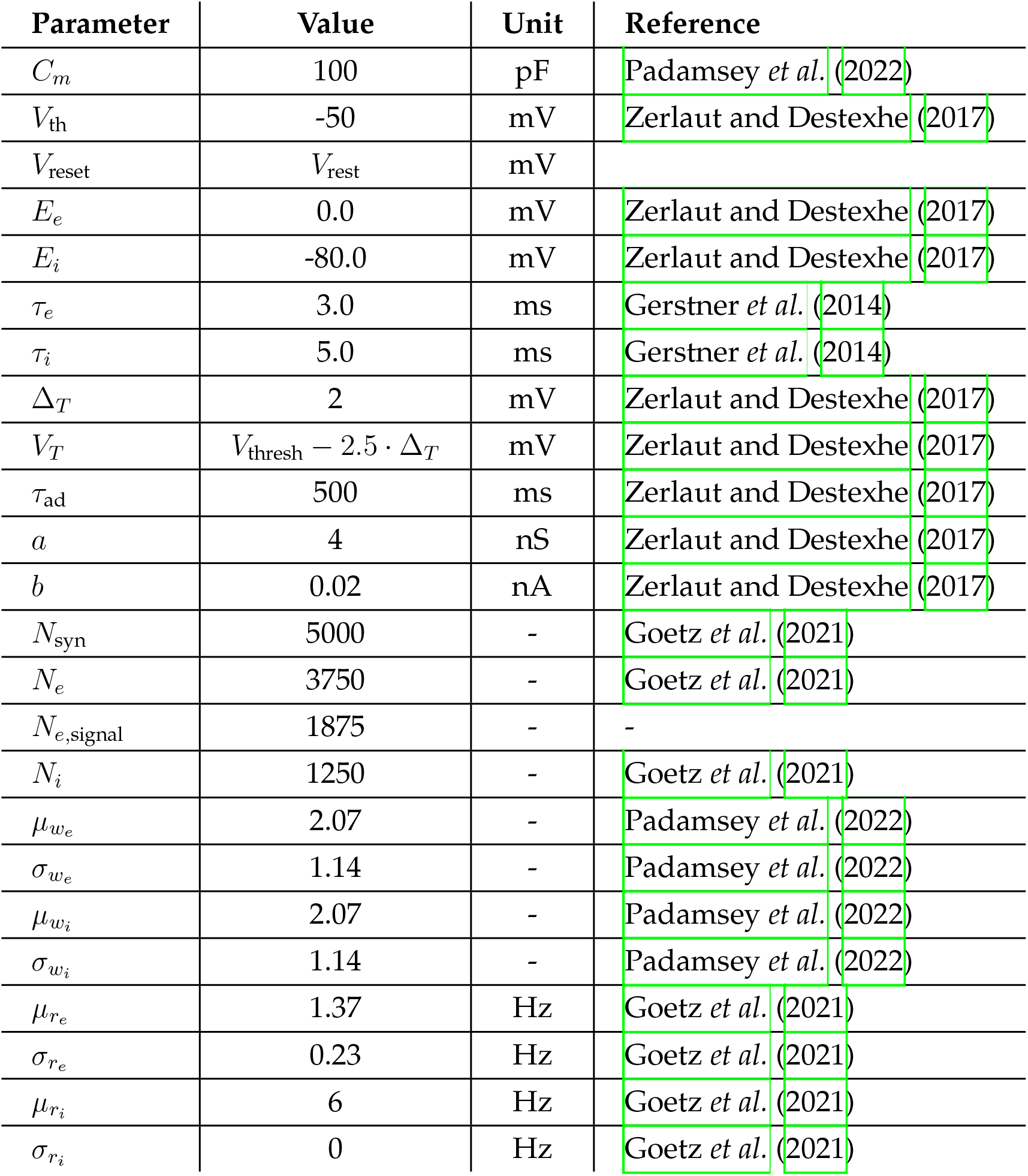
Parameters for the LIF and AdExp neuron model and oriented gratings synaptic inputs.

#### Model switching ON-OFF states as synaptic input

Following the approach of Zeldenrust *et al*. (2017), we modelled the stimulus as a hidden binary state (Markov process) switching stochastically between preferred (ON) and non-preferred (OFF) conditions. The transition from OFF to ON occurred at a rate *r*_ON_, and from ON to OFF at *r*_OFF_. The resulting switching time constant of the hidden state is given by:

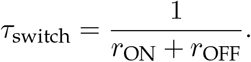

In line with experimental observations (Zeldenrust et al., 2024), excitatory and inhibitory neurons exhibit distinct operational timescales — ranging from a few Hz in excitatory cells to over 20 Hz in inhibitory neurons. Accordingly, the switching dynamics of the inputs were adapted: we used *τ*_switch_ = 50 ms for excitatory and 250 ms for inhibitory neurons to match their respective membrane integration properties.

Synaptic inputs were modelled analogously to earlier sections: excitatory and inhibitory synaptic weights were drawn from a log-normal distribution (Fig. S11A), excitatory presynaptic firing rates from a normal distribution (Fig. S11B), and inhibitory rates were set to a uniform distribution. Stimulus encoding was implemented via a matching scheme: during ON epochs, strong synapses were paired with high firing rates, whereas during OFF epochs, this pairing was inverted (Fig. S11C,D):

- For the preferred stimulus (ON) large synapses are assigned to high firing rates, small synapses with low firing rates.
- For the non-preferred stimulus (OFF) large synapses fire with low firing rates, small synapses with high firing rates.

The corresponding firing rates were translated into spiking input with an inhomogeneous Poisson process (a time-varying rate code).

For subsequent analyses, parameter grids spanning *±*3 standard deviations in the nine-dimensional principal component space (see Fig. S14) were simulated separately for excitatory and inhibitory neurons.

**Table 2:** Parameters for the LIF and AdExp neuron model and switching ON-OFF synaptic inputs.

| Parameter | Value | Unit | Reference |
| --- | --- | --- | --- |
| $R_m$ | 100 | mΩ | Zeldenrust *et al.* (2024) |
| $C_m$ | 50 | pF | Zeldenrust *et al.* (2024) |
| $V_{\text{rest}}$ | -70 | mV | Zeldenrust *et al.* (2024) |
| $V_{\text{th}}$ | -63 | mV | Zeldenrust *et al.* (2024) |
| $V_{\text{reset}}$ | $V_{\text{rest}}$ | mV | |
| $E_e$ | 0.0 | mV | Zerlaut and Destexhe (2017) |
| $E_i$ | -80.0 | mV | Zerlaut and Destexhe (2017) |
| $\tau_e$ | 3.0 | ms | Gerstner *et al.* (2014) |
| $\tau_i$ | 5.0 | ms | Gerstner *et al.* (2014) |
| $\Delta_T$ | 1 | mV | Zeldenrust *et al.* (2024) |
| $V_T$ | $V_{\text{thresh}} - 2.5 \cdot \Delta_T$ | mV | Zerlaut and Destexhe (2017) |
| $\tau_{\text{ad}}$ | 10 | ms | Zeldenrust *et al.* (2024) |
| $a$ | 4 | nS | Zeldenrust *et al.* (2024) |
| $b$ | 0.0805 | nA | Zeldenrust *et al.* (2024) |
| $N_{\text{syn}}$ | 1333 | - | Goetz *et al.* (2021) |
| $N_e$ | 1000 | - | Zeldenrust *et al.* (2024) |
| $N_{e,\text{signal}}$ | 500 | - | - |
| $N_i$ | 333 | - | - |
| $r_{\text{ON}_e}$ | 1.3 | - | Zeldenrust *et al.* (2024) |
| $r_{\text{OFF}_e}$ | 2.7 | - | Zeldenrust *et al.* (2024) |
| $r_{\text{ON}_i}$ | 6.7 | - | Zeldenrust *et al.* (2024) |
| $r_{\text{OFF}_i}$ | 13.3 | - | Zeldenrust *et al.* (2024) |
| $\mu_{w_e}$ | 2.07 | - | Padamsey *et al.* (2022) |
| $\sigma_{w_e}$ | 1.14 | - | Padamsey *et al.* (2022) |
| $\mu_{w_i}$ | 2.07 | - | Padamsey *et al.* (2022) |
| $\sigma_{w_i}$ | 1.14 | - | Padamsey *et al.* (2022) |
| $\mu_{r_e}$ | 1.37 | Hz | Goetz *et al.* (2021) |
| $\sigma_{r_e}$ | 0.23 | Hz | Goetz *et al.* (2021) |
| $\mu_{r_i}$ | 6 | Hz | Goetz *et al.* (2021) |
| $\sigma_{r_i}$ | 0 | Hz | Goetz *et al.* (2021) |

#### Information measures

Many information measures have the caveats that they can only be applied to specific situations with binary sensory input. We calculate four different information measures:

- mutual information *MI* between in-& output,
- transfer entropy *TE* between in-& output with one time bin as lag,
- coding efficiency of a spike train *CE*,
- tuning curve,
- orientation selectivity index *OSI*,
- coefficient of variation of neuronal activity *CV*_ISI_,
- coefficient of variation of subthreshold membrane variability 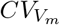,

and relate them to the energy consumption.

The mutual information (*MI*) measures the overlap between the information content of two (sub)systems Shannon (1948). The mutual information between a given stimulus *s* and the neuronal response/firing rate *r* is denoted as *MI*(*r*|*s*) (for continous time series (Zeldenrust *et al*., 2017)) and given by:

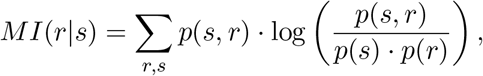

where *p*(*s, r*) is the joint probability describing the likelihood of observing a particular combination of *s* and *r* concomitantly, *p*(*s*) the stimulus probability of observing particular stimulus *s*, and *p*(*r*) the neuronal response probability of observing particular neuronal response *r*. It is a symmetric measure where no distinction between cause and effect can be made.

By introducing transition probabilities into mutual information dynamical and directional information transport can be quantified. The resulting measures is called transfer entropy (*TE*). This asymmetrical measure estimates causally transmitted information with a directed exchange of information to distinguish effectively driving and responding signal with specific time lag (Schreiber, 2000). The information flow from a given stimulus *s* to the neuronal response/firing rate *r* considering the the past stimulus *s*′ with time lag *τ* is denoted as the transfer entropy *TE*(*r*|*s, τ*) and is given by:

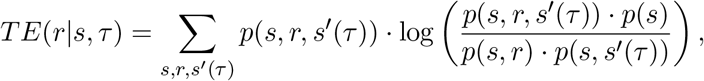

where *p*(*s, r, s*′ (*τ*)) is the joint probability describing the likelihood of observing a particular combination of current stimulus *s*, neuronal response *r*, and past stimulus *s*′ delayed by a time lag *τ* concomitantly, *p*(*s*) is the stimulus probability of observing particular stimulus *s, p*(*s, r*) is the joint probability describing the likelihood of observing a particular combination of *s* and *r* concomitantly without considering past stimuli, and *p*(*s, s*′ (*τ*)) is the joint probability of the current stimulus *s* and the past stimulus *s*′ describing the relationship between the current and past states of a stimulus.

To estimate the mutual information and transfer entropy the spike trains were discretized in time and frequency space. The bin sizes were optimized by maximizing the information content for different combinations of time and frequency bins (optimal combinations: bin time_*MI*_ = 1200 ms and bins rate_*MI*_ = 150 and bin time_*TE*_ = 294 ms and bins rate_*TE*_ = 58. The time lag to estimate the transfer entropy was one bin size (*τ* = 294 ms), see Fig. S5).

The coding efficiency of a spike train *CE* describes how much information is transmitted per spike. It is calculated by dividing the information content of a spike train (either measured by *MI* denoted as *CE*_*MI*_ or by *TE* denoted as *CE*_*TE*_) by the number of spikes in the train represented by the firing rate *r*_post_:

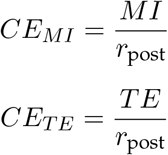

The tuning curve describes the response of the neuron to presented stimuli. The neuronal response is estimated by the firing rate of the neuron during each stimulus condition presentation.

The orientation selectivity index describes how selective a neuron responds to a stimulus. It compares the response of a neuron to its preferred orientation *r*_pref_ to its non-preferred (equivalently orthogonal, 90° shifted grating relative to preferred orientation) orientation response *r*_ortho_:

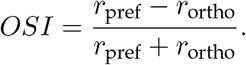

An *OSI* of 1 describes a selective neuron which has exclusive/strong response for preferred and no response to non-preferred stimuli. Conversely, an *OSI* of 0 describes a neuron responding to all stimuli equally making it non-selective for this group of stimuli.

The coefficient of variation of neuronal activity *CV*_ISI_ is a measure for variability of the inter-spike intervals of a spike train:

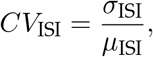

where *σ*_ISI_ is the standard deviation of inter-spike intervals and *µ*_ISI_ is the mean inters-spike interval duration.

The coefficient of variation of subthreshold membrane variability 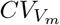 describes how variable the subthreshold membrane potential *V*_*m*_ is:

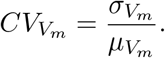

#### Zeldenrust fraction of transferred information

To quantify the information transferred by the neuron for switching ON-OFF stimulation, we use the fraction of transferred information (*FI*) the ratio between the mutual information between postsynaptic spike train (*MI*_spike train_) and hidden state and the mutual information between synaptic input and hidden state (*MI*_input_):

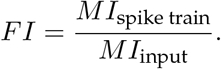

This quantifies the amount of information kept during the spike generation process.

The used information measures of our model assume rate-based coding, whereas real neurons likely employ a combination of rate, phase, and temporal coding strategies.

#### Energy budget

With their landmark study Attwell and Laughlin (2001) gave a simple estimate of the total energy of grey matter neurons as the sum of different subcellular processes. It was later updated by Howarth *et al*. (2012) and biologically validated by Yu *et al*. (2017) who compared expected and measured glucose consumption. Following the aforementioned studies, we assume the hydrolysis of ATP as the main energy provider and thus ATP as the main energy carrier (“energy currency of the cell”). For all energy budgets we calculate how much ATP a neuron consumes per second.

The total energy demand of neurons can be split into a non-signalling (house-keeping and maintaining a resting potential) and signalling component being activity/frequency dependent (action potential generation and conductance, synaptic transmission process, glutamate/GABA recycling, presynaptic Ca^2+^ entry). The total energy demand splits into six main subprocesses:

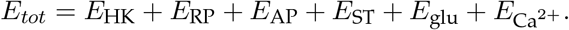

The different components can be estimated separately and vary due to the type and amount of molecules and ions involved and also depend on the cell morphology.

##### House keeping energy budget

Cells have a basic demand of energy to maintain their fundamental functions and shape such as synthesis and turnover of proteins, phospholipids and other macromolecules, microtubule cytoskeletal rearrangements and turnover, axoplasmic transport by actin tread milling, and mitochondrial proton leak (Engl and Attwell, 2015) (synaptogenesis (Balasubramanian, 2021)). These energy demands do not dependent on the activity of the neuron and thus are also consumed when the neuron is silent. All these house-keeping processes are summed in the term *E*_HK_ and represent how many hydrolyses of ATP molecules per second a neuron demands.

We chose to keep the house-keeping energy demand constant for all activity regimes following Yu et al. (2017). Attwell and Laughlin (2001) and Howarth *et al*. (2012) calculated it as 1/3 of the signalling energy and others report 1/5 (Hyder *et al*., 2013). The effect of this assumption increases with firing rate. Because our results suggest medium-low firing-rates being the energetically most efficient ones, we argue that the effect of this assumption has only minor effect on the results. But for investigating neurons with high firing rates the activity-dependence of the house-keeping energy demand should be considered further. Furthermore, Balasubramanian (2021) report that 10 % of total energy remains unused under normal operation and serves as energy reserviour to buffer periods of high energetic demands.

##### Resting potential energy demand

To maintain the resting potential, a state maintained far from thermodynamic equilibrium, a neuron continuously needs to pump sodium out of the cell (and potassium into the cell) due to the leakiness of the cellular membrane to ions. Pumping sodium against the electrical (charge difference) and chemical (concentration difference) gradient involves the sodium-potassium ion pump which extrudes 3 Na^+^ and imports 2 K^+^ under the consumption of 1 molecule of ATP. Assuming that the cell membrane at rest is (only) permeable to Na^+^ and K^+^ and all charge fluxes through these leaky conductances are compensated by the sodium-potassium pump with given stoichiometry and ATP demand (in opposite direction) (no net charge flux, balanced sodium and potassium conductances), Attwell and Laughlin (2001) derive Formula (4) in their paper to estimate the energy demand *E*_RP_ to maintain the membrane at a specific resting potential *V*_RP_:

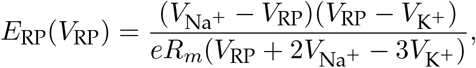

where *V*_Na_+ is the sodium reversal/Nernst potential, *V*_K_+ the potassium reversal/Nernst potential, *e* the elementary charge, and *R*_*m*_ the membrane resistance. The somatic estimate of input resistance derived from our experimental data slightly overestimates the true

membrane resistance relevant for calculating the resting potential energy, by approximately 9%. To assess the overestimation of membrane resistance from somatic input resistance measurements, we performed simulations using a detailed compartmental model of a layer 2/3 pyramidal neuron (Goetz et al., 2021). A current of 0.05 nA was injected into the soma, resulting in a voltage deflection of 7.331 mV and an input resistance (*R*_in_) of 146.62 M”. Given the model’s total surface area (*A*_neuron_ = 0.000111 cm^2^), this corresponded to an estimated membrane resistance (*R*_*m*_ = *R*_in_ · *A*_neuron_) of 16.3 k·Ω ·cm^2^. In contrast, the model’s true specific membrane resistance was 15 kΩ·cm^2^, indicating that somatic input resistance overestimates the true membrane resistance by approximately 9%. This bias is likely even larger for the larger neurons considered by Attwell and Laughlin (2001). We therefore applied a correction factor based on this estimate when calculating energy budgets from experimental measurements.

We are aware, that the above equation to estimate the resting energy represents a conservative estimate of energy savings as drastically changing the resting potential violates some of the original assumptions made by Attwell and Laughlin (2001). In particular, this equation assumes that the resting potential is governed by the ratio of just one conductance for each of Na^+^ and K^+^. A higher resting potential therefore implies a higher Na^+^ conductance which actually increases ATP cost (Fig S8). In reality, the membrane potential is maintained by a larger number of ionic conductances. Hyperpolarization-activated cyclic nucleotide-gated (HCN) channels depolarize the resting potential, contribute to resting potential stability and dendritic integration (Wahl-Schott and Biel, 2008). By incorporating HCN-mediated currents into the energy budget, we refine estimates of resting potential maintenance. The somato-dendritic gradient of HCN channels also influences temporal integration, potentially tuning synaptic efficacies and response selectivity (Zhuchkova et *al*., 2013). The *I*_*h*_ might also be adapted during food-restriction helping to save energy (more degrees of freedom). Thus, for our energy calculations we extent the original formula derived by (Attwell and Laughlin, 2001) by an additional *I*_*h*_ resulting in

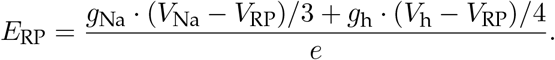

The full derivation with the basic assumptions is presented in the supplements.

##### Action potential energy budget

Firing an action potential and propagating it to along the whole axon (of a neuron to the dendritic terminals) involves a depolarization (sodium influx, rising phase) followed by a repolarization or even a hyperpolarization (potassium efflux, falling phase) of the neuronal membrane. The re- and hyperpolarization do not happen spontaneously because intracellular sodium has to be extruded against the electrochemical gradient. Thus, the energy demand of an action potential can be calculated by the total sodium charge influx into the cell during depolarization (due to an action potential being fired), which has to be extruded by the sodium-potassium pump (with stoichiometry 3 Na^+^ per ATP) during repolarization under the consumption of ATP (Attwell and Laughlin, 2001):

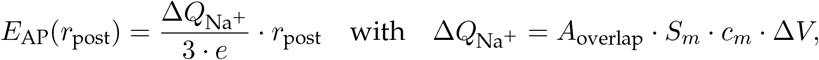

where *E*_AP_ is the energy demand for action potential firing at a mean firing rate *r*_post_, Δ*Q*_Na_+ is the sodium charge influx which has to be extruded by the sodium-potassium pump, *e* is the elementary charge, *A*_overlap_ is the overlap factor between sodium and potassium currents, *S*_*m*_ is the surface area of the neuron, *c*_*m*_ is the specific membrane capacitance, and Δ*V* being the difference between the spike peak voltage *V*_*peak*_ and the threshold voltage *V*_th_:

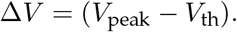

The surface area of a neuron is estimated as the sum of the surface area of the axon, soma, and the dendritic tree (Attwell and Laughlin, 2001):

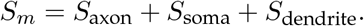

The surface area of the axons is calculated by:

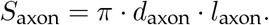

The surface area of the soma is calculated by:

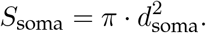

The surface area of the dendritic tree is calculated by:

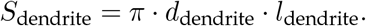

The theoretical minimum charge needed to depolarize the membrane from *V*_th_ to *V*_*peak*_ (amplitude of the action potential) is Δ*Q*_Na_+_,min_ = *S*_*m*_ · *c*_*m*_ · Δ*V* . In biological neurons, however, sodium influx and potassium efflux overlap temporally due to channel kinetics, increasing the effective sodium charge influx (Sengupta *et al*., 2010). This excess cost by increased sodium charge influx is commonly described by the overlap ratio *A*_overlap_ and describes the energy-efficiency of action potentials.

Attwell and Laughlin (2001) calculated the contribution of action potentials to their energy budget based on measurements from the squid giant axon, where temporal sodium–potassium current overlap results in an energy expenditure approximately four times the theoretical minimum (Hodgkin and Huxley, 1952; Hodgkin, 1975) However, the squid giant axon is an energetically inefficient reflex neuron compared to most mammalian neurons. Alle et al. (2009) (for rat hippocampal non-myelinated mossy fibers) and Carter and Bean (2009) (for mouse pyramidal cells in hippocampal CA1 and neocortex) showed that action potentials in mammalian neurons are much more energy-efficient by measuring the overlap between the rapidly decaying sodium current and delayed-onset potassium current waveforms. The authors showed that action potentials in mammalian excitatory axons and somata cost only 1.3 instead of 4 times the theoretical minimum. These findings led to a revised and lower estimate of spike-related energy consumption in mammalian cortex (Howarth *et al*., 2012).

Carter and Bean (2009) also provided experimental evidence that the amount of sodium entry during action potential generation varies considerably among different species and cell types (Carter and Bean (2009), Fig. 4). An increased sodium influx due to an incomplete sodium inactivation decreases energy efficiency but enables fast spiking (narrower APs) as a functional benefit (Carter and Bean, 2009). Thus, it is important to consider the type of neurons because they trade-off energy efficiency and function differently.

Alternative approaches for estimating the energetic cost of action potential generation have also been proposed (Yi *et al*., 2016). Rather than quantifying ion flux directly, these methods infer energy demand from the electrical power dissipated by ionic conductances during spiking, offering a complementary perspective to ion-count-based estimates.

It is important to note that the membrane capacitance *C*_*m*_ reported in Padamsey *et al*. (2022) reflects the somatic capacitance only. While this value is appropriate for simulating integrative properties in single-compartment neuron models (LIF or AdExp), it underestimates the total energy cost of action potential generation, which also involves the axon and dendritic arbor. To accurately capture the full energetic demand, we compute the total membrane capacitance by multiplying the specific membrane capacitance (*c*_*m*_) with the complete surface area (*S*_*m*_) of a pyramidal neuron.

We additionally assessed the energetic contribution of adaptation currents. The mean adaptation current increases with firing rate and with the adaptation time constant (Fig. S7A). Within the firing-rate range relevant for optimal coding in our study (up to ∼ 5 Hz), adaptation currents contribute no more than 13 % of the mean excitatory synaptic current (Fig. S7B). To examine the ionic composition and energetic cost of adaptation currents during spike generation, we stimulated a detailed compartmental model (Goetz et al., 2021) with brief step current injections (15 ms duration), once with adaptation currents (*I*_*M*_ and *I*_*K*,Ca_) enabled and once with these currents disabled (Fig. S7C). Tracking ionic fluxes revealed that the presence of adaptation currents slightly reduces the total ionic load per action potential, resulting in a lower energetic cost per spike (Fig.S7D). Given their minor contribution in the low firing-rate regime studied here, adaptation currents were not explicitly included in the energy budget. However, their energetic impact may become relevant at higher firing rates and under stronger adaptation.

##### Synaptic transmission energy budget

The release of neurotransmitter from the presynaptic terminal and their binding to postsynaptic (ionotropic) receptors results in an influx of Na^+^ and Ca^2+^. We calculated the energy demand of the postsynaptic cell to re-establish the resting ion gradients due to presynaptic activity as:

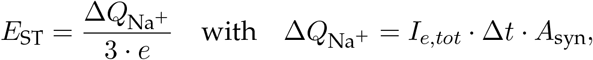

where *E*_ST_ is the energy demand to restore the resting membrane potential due to the “effective” (post)synaptic sodium charge influx Δ*Q*_Na_+ which has to be extruded by the sodium-potassium pump. The effective sodium charge influx Δ*Q*_Na_+ is the product of the total excitatory synaptic current *I*_*e,tot*_ entering the cell in time interval Δ*t* (energy measures per second demands Δ*t* = 1 s), and the proportionality factor *A*_syn_ which accounts for the different stoichiometry of sodium and calcium (potassium) influx as well as inhibitory chloride influxes (which have a much smaller electrochemical gradient at inhibitory synapses and thus consume way less energy than sodium at excitatory synapses (Attwell and Laughlin, 2001; Howarth *et al*., 2012). Additionally, Attwell and Laughlin (2001) treat all neurons as glutamatergic since they outnumber inhibitory neurons by 9:1 and 90 % of synapses release glutamate (Braitenberg and Schü z, 1998).

We derive *A*_syn_ as follows:

Assuming that the (measured/simulated) excitatory synaptic current consists only of sodium ions the energy demand to reverse the influx of sodium charge Δ*Q*_Na_+ = *I*_*e,tot*_ · Δ*t* (per second Δ*t* = 1 s) is:

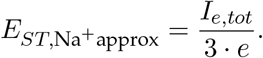

To obtain biophysically grounded estimates of synaptic ionic currents, we leveraged a detailed multi-compartment model of a layer 2/3 pyramidal neuron (Goetz *et al*., 2021). Although this model was originally parameterized for rat cortex, comparative analyses across mammalian species indicate that key electrophysiological and metabolic scaling relationships are largely conserved, with humans representing the primary outlier in allometric scaling (Beaulieu-Laroche et al., 2021). We therefore use this model as a representative mammalian reference to estimate the relative contributions of ionic currents during synaptic transmission. Using this framework, we quantified the mean dendritic ionic currents associated with excitatory synaptic input, yielding the following average synaptic current components:

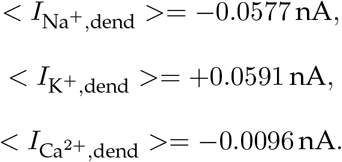

The ratio of Na^+^-to Ca^2+^-currents is approximately 6:1. Thus, the Ca^2+^-current accounts for 1*/*7 of the total current (*I*_*e,tot*_). Splitting the excitatory synaptic current into its relative contributions 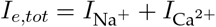:

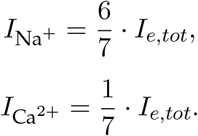

The ATP-demand to extrude Ca^2+^ is much different to extrude Na^+^. One molecule of ATP extrudes 3 Na^+^ or 1 Ca^2+^. Accounting for the double charge of Ca^2+^ and the ATP-stoichiometry the energy consumption of the contributions are:

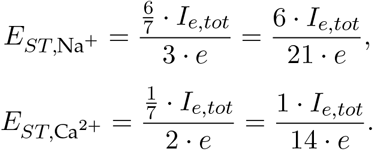

In sum this results in:

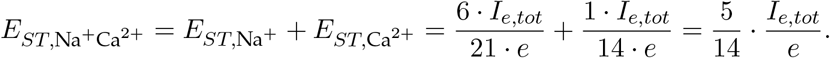

Setting this measure in relation to *E*_*ST*,Na_+_approx_ we receive a factor *A*_syn_ which corrects the energy estimate of the non-specific excitatory synaptic current approximation (of the excitatory synaptic currents only consisting of Na^+^) by accounting for different stoichiometries of Na^+^ and Ca^2+^:

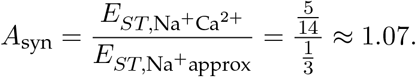

The mean ionic currents of the compartmental model vary for different stimuli presentations in a range from 2:11 for preferred till 1:8 for non-preferred. Calculating *A*_syn_ analogously to before these ratios results in a range of *A*_syn_ of 1.055 for non-preferred until 1.08 for preferred stimuli, thus, not varying a lot. To account also for energetic demands of inhibitory synapses we set this correction factor to *A*_syn_ ≈ 1.1.

Attwell and Laughlin (2001) already mention the estimation of synaptic transmission energy from postsynaptic ion fluxes but also that few experimental data are available. But the postsynaptic ionic currents are easily accessible (inate output variable) of our neuronal modeling. Attwell and Laughlin (2001) calculated the energy budget of the synaptic transmission process by the presynaptic activity:

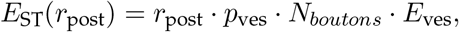

where *p*_ves_ is the release probability for glutamate vesicles, *N*_*boutons*_ the number of downstream synaptic contacts to following cells, and *E*_ves_ is the energy needed to reverse the sodium ion influx through NMDA and non-NMDA ionotropic receptors and metabotropic receptors (G-protein signalling) due to one vesicle of glutamate of around 137 000 ATP molecules per vesicle (Attwell and Laughlin, 2001). This calculation assumes a linear increase of synaptic transmission energy demand with the firing rate and that that the pre- and postsynaptic firing rates always behave equally. This assumption is based on the idea that the release of presynaptic glutamate results in the opening of an equal amount of postsynaptic receptors and thus constant postsynaptic influx of ions and metabolic consequences. But this assumption of symmetry in pre- and postsynaptic firing rate/energy use neglects the postsynaptic non-linear integration process and does not capture synaptic scaling. These estimations of ATP consumption due to synaptic transmission make the simplifying assumption used in Attwell and Laughlin (2001) that neurons in a population are recurrently connected so that pre- and post-synaptic firing rates are equal. For sensory systems with a more hierarchical organisation (Van Essen and Maunsell, 1983), it is likely that these two rates will differ, so we relax this assumption here.

##### Neurotransmitter recycling energy budget

Excitatory synaptic signalling involves mainly the presynaptic release of glutamate and its binding to glutamate receptors at the postsynaptic cell. The released glutamate is resumed by the presynaptic cell, metabolically recycled/reprocessed, repacked in vesicles, and transported to the dendritic terminals for further signalling (Li and Sheng, 2022). We calculated the energy demand of the cell for neurotransmitter recycling (mainly glutamate) *E*_glu_ as:

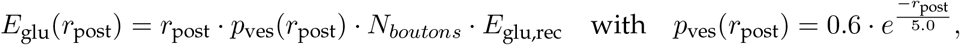

where *p*_ves_ is the release probability for glutamate vesicles depending on the neuronal firing rate *r*_post_, *N*_*boutons*_ the number of downstream synaptic contacts/boutons to neurons, and *E*_glu,rec_ the energy to recycle one vesicle of glutamate. One vesicle contains approximately 4000 glutamate molecules (Riveros *et al*., 1986) and to recycle one glutamate molecule 3.67 ATP molecules are needed (Yu *et al*., 2017). The presynaptic release probability *p*_ves_ is assumed to decay exponentially as a function of presynaptic firing rate (*r*_post_) Attwell and Laughlin (2001); Howarth *et al. (2012);* Yu *et al. (2017)* due to the depletion of neurotransmitter during prolonged stimulation (Del Castillo and Katz, 1954). The steady-state release probability is an exponentially decreasing function of rate (Tsodyks and Markram, 1997) for a broad class of presynaptic firing patterns (Bird and Richardson, 2018). Attwell and Laughlin (2001); Howarth et al. (2012) used the simplification of a constant value of *p*_ves_ = 0.25 to estimate the release probability for a mean activity of *r*_post_ = 4 Hz. But our simulations involve a broad range of different firing rates. Thus, we account for the (quick) exponential decay of presynaptic release probability with a time constant of the synaptic depression of 200 ms (Tsodyks *et al*., 1998). Moreover, the presynaptic release probability has been found to adapt to energy levels by regulating the speed of synaptic vesicle cycle (Yuan *et al*., 2018). This is in line with findings of activity-enhanced presynaptic energy production (Li and Sheng, 2022).

##### Presynaptic calcium extrusion energy budget

The synaptic transmission process includes an influx of calcium which triggers, if enough calcium accumulates in a bouton, the release of a neurotransmitter vesicle. This intracellular calcium has to be extruded against the electrochemical gradient, which consumes energy. Another consumer of energy is the vesicular release (endocytosis) in form of a fusion of vesicle and cell membrane. Similar to the glutamate release energy budget, also the cost for calcium extrusion is activity-dependent. Analogously, we calculated the energy demand of the cell for calcium extrusion *E*_Ca_2+ as:

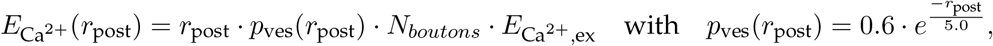

where *E*_Ca_2+_,ex_ is the energy needed to extrude the calcium influx of a single presynaptic action potential to release one neurotransmitter vesicle. For the release of one vesicle 12 000 Ca^2+^ ions are needed and it takes 1 ATP molecule to extrude 1 Ca^2+^ ion (Attwell and Laughlin, 2001).

Additional minor presynaptic energy contributors are synapse assembly, their maintainance by cargo transport (Li and Sheng, 2022).

##### Minor energy consumption of inhibitory currents

Neuronal energy consumption depends on morphology (size of membrane area), synaptic connectivity, the precise combination and densities of ionic conductances, their type, function and neurotransmitter. While we focused on Na^+^, K^+^, and Ca^2+^ gradients, other ions (H^+^, Cl^−^, and 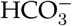) also contribute to energy costs (Attwell and Laughlin, 2001). Inhibition, while metabolically less demanding and often neglected (around 1 % of the total energy demand (Attwell and Laughlin, 2001; Howarth et al., 2012)), plays a critical role in network function (Buzsaki *et al*., 2007; Raimondo *et al*., 2017) and was incorporated into our model through a synaptic proportionality factor *A*_syn_ derived from a compartmental model (Goetz *et al*., 2021).

##### Minor energy consumption of adaptation currents

In addition, the energy cost of adaptation currents may merit consideration. As shown by Yi *et al*. (2016), spike-frequency adaptation mediated by Ca^2+^-activated (*I*_AHP_) and K^+^-activated (*I*_M_) currents nearly doubles the energy cost of action potentials under strong adaptation. These slow hyperpolarizing K^+^ currents increase the sodium influx required to reach threshold, thereby enhancing the overlap between inward sodium and outward potassium currents and reducing action potential efficiency. To account for this additional cost, it may be necessary to introduce either a frequency-dependent adjustment of the overlap factor (*A*_overlap_) or an explicit energy term linked to the adaptation current (*w*_ad_), which remains to be quantified. In our simulations, the mean adaptation current increased with firing rate (Fig. S7A) but remained below 14 % of the mean excitatory current across physiologically relevant firing rates (Fig. S7B). Interestingly, disabling adaptation currents (*I*_M_ and *I*_K,Ca_) in a detailed compartmental model (Goetz *et al*., 2021) increased the energetic cost of a single spike by 5%, due to greater sodium and calcium influx(Fig. S7C-D).

In total, the reversal of ion-fluxes consumes the largest amount of energy with the synaptic transmission processes being the main contributor.

**Table 3:** Neuronal parameters to calculate the energy budget of a layer 2/3 pyramidal excitatory neuron of rats with cellular anatomies, specific electrical properties, and natural constants. To calculate energy demands of human neurons simply multiply *l*_axon_ and *N*_*boutons*_ by 1.89 to account for their larger size and different synaptic densities (higher metabolism) (Yu *et al*., 2017).

| Parameter | Value | Unit | Reference |
| --- | --- | --- | --- |
| $E_{HK}$ | $5.38 \cdot 10^8$ | ATP/s | Yu et al. (2017) |
| $V_{Na^+}$ | 50 | mV | Attwell and Laughlin (2001) |
| $V_{K^+}$ | -100 | mV | Attwell and Laughlin (2001) |
| $e$ | $1.6 \cdot 10^{-19}$ | C | - |
| $A_{overlap}$ | 1.24 | - | Carter and Bean (2009) |
| $d_{axon}$ | 0.3 | $\mu m$ | Braitenberg and Schüz (1998) |
| $l_{axon}$ | 4 | cm | Braitenberg and Schüz (1998) |
| $d_{soma}$ | 25 | $\mu m$ | Attwell and Laughlin (2001) |
| $d_{dendrite}$ | $3 \cdot d_{axon}$ | $\mu m$ | Braitenberg and Schüz (1998) |
| $l_{dendrite}$ | $l_{axon}/9$ | cm | Braitenberg and Schüz (1998) |
| $c_m$ | 0.01 | F/m<sup>2</sup> | Attwell and Laughlin (2001) |
| $V_{peak}$ | 25 | mV | Carter and Bean (2009) |
| $V_{th}$ | -50 | mV | Zerlaut and Destexhe (2017) |
| $A_{syn}$ | 1.1 | - | |
| $N_{boutons}$ | 8000 | - | Braitenberg and Schüz (1998) |
| $E_{glu,rec}$ | $1.468 \cdot 10^4$ | ATP/s | Yu et al. (2017) |
| $E_{Ca^{2+},ex}$ | $1.2 \cdot 10^4$ | ATP/s | Attwell and Laughlin (2001) |

## Supporting information

## Frequently used abbreviations

AdExp: Adaptive exponential integrate-and-fire neuron model
ATP: Adenosine triphosphate
CE: Coding efficiency
CTR: Control condition; normally fed mice
CV: Coefficient of variation
FR: Food-restricted condition; mice subjected to 2 weeks of dietary restriction
FWHM: Full width at half maximum
HCN: Hyperpolarization-activated cyclic nucleotide-gated (HCN) channels
ISI: Inter-spike interval
LIF: Leaky integrate-and-fire neuron model
mEPSC: Miniature excitatory postsynaptic current
MI: Mutual information
OSI: Orientation selectivity index
SBI: simulation based inference
TE: Transfer entropy

---

**Figure S1:**
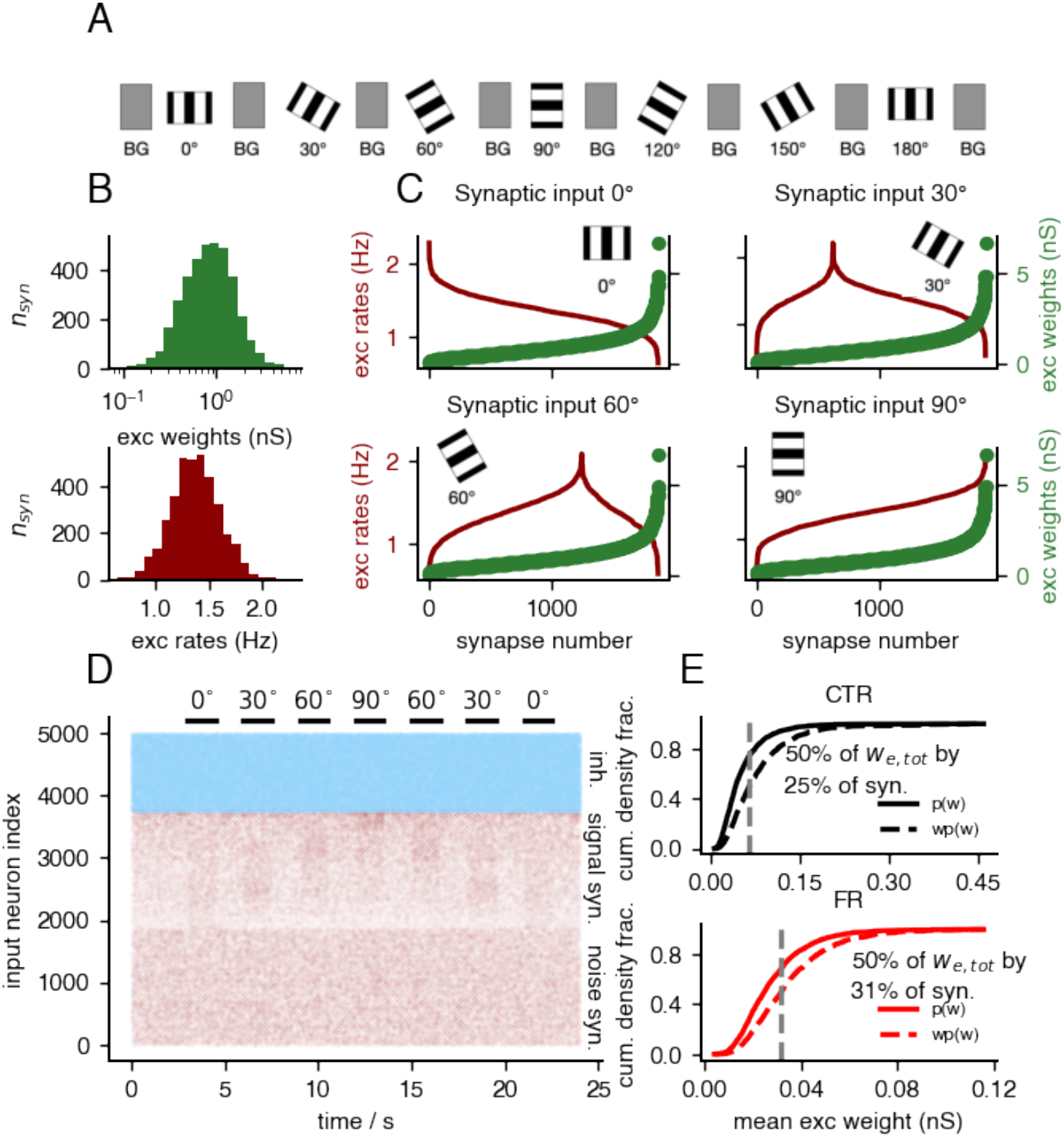
Design of *in vivo*-like excitatory and inhibitory synaptic input to a layer 2/3 pyramidal neuron in primary visual cortex with orientation variation stimulus. **A**, Oriented gratings are present for 1.5 s with 1 s intermittent background firing (BG) and the orientation values 0°, 30°, 60°, 90°, 120°, 150°, 180° as done by Padamsey et al. (2022). **B – Top**, Excitatory synaptic weights follow a log-normal distribution 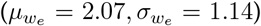. **B – Bottom**, Excitatory presynaptic firing rates are normally distributed 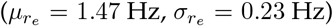. **C**, The seven orientations are assigned synaptic inputs based on the matching scheme, visualized as cityscape histograms: presynaptic firing rates (dark red) as a function of synaptic strength (green). The preferred stimulus (90°) elicits the strongest total synaptic drive, whereas non-preferred orientations (0°, 180°) exhibit the weakest. Intermediate orientations fall between these extremes. **D**, Raster plot of synaptic input spike times, distinguishing inhibitory (light blue), excitatory signal and excitatory noise (red as a function of the synaptic strength). **E – Top**, Cumulative density function of the experimentally observed mEPSC distribution as a proxy for synaptic strength in control cells (Song et al., 2005. **E– Bottom**, The same for food-restricted cells, showing a disproportionate reduction in the strongest synapses under food restriction reduces the effect of the strongest synapses.

## Broadness of tuning curve

The strong broadening of tuning curves under food restriction suggests that synaptic weights undergo non-multiplicative downscaling. Strong synapses are preferentially weakened, reducing overall energy expenditure but reducing the selectivity of the neuron. But, it is unclear if the *ex vivo* measured synaptic weight distribution represent the *in vivo* scaling behavior. The broadness of weight distribution affects the neuronal response and the width of tuning curve in opposite way: narrow weight distribution lead to broad tuning curves because only few synapses are coding for the preferred stimulus (Fig. 5A–C). The number of signaling synapses significantly influences the sharpness of tuning curves. In Fig. S2A, we show that a higher proportion of signaling synapses leads to a more pronounced tuning curve. This effect arises from the log-normal distribution of synaptic strengths in our input model. For the preferred orientation (90°), strong synapses are paired with high firing rates, resulting in a strong synaptic drive. When the fraction of signaling synapses increases, the contribution of strong synapses becomes more dominant relative to weaker ones, further enhancing the response to the preferred stimulus. The resulting tuning curves of the 50% signaling synapse ratio matched the experimental data best.

**Figure S2:**
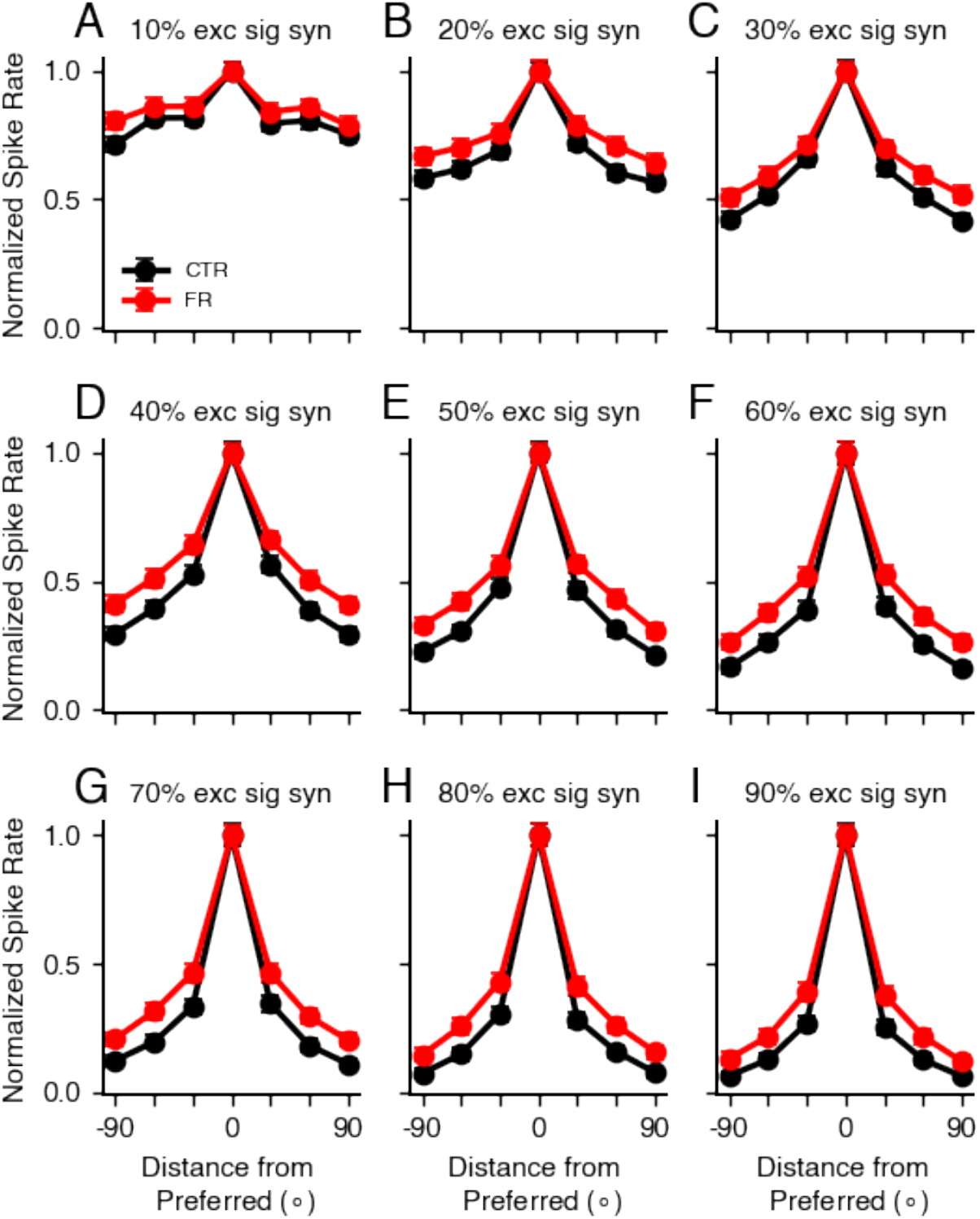
A higher signaling synapse ratio increases tuning curve sharpness. Using an AdExp model with orientation selectivity task (Fig. S1) shows that tuning curves for different ratios of signaling synapses (**A**, 10% to **I**, 90%) show increased peakiness with a higher proportion of signaling synapses. This effect arises from the log-normal distribution of synaptic strengths: for the preferred orientation (90°), strong synapses are paired with high firing rates, leading to a stronger synaptic drive. As the proportion of signaling synapses increases, the relative contribution of strong synapses becomes more dominant, enhancing the selectivity of the neuronal response. The tuning curves with a excitatory signaling synapse ratio of 50% matched the experimental tuning curves (Padamsey *et al*., 2022) best.

## Adaptation and spike-generation non-linearities shape tuning curve broadening

At experimentally measured levels of intrinsic membrane noise, a generic leaky integrate- and-fire (LIF) neuron fails to reproduce the tuning curve broadening observed under food restriction (Fig. S3A) in simulations with oriented grating input (Fig. S1). Introducing an adaptation current does not qualitatively alter this behavior (Fig. S3B), indicating that adaptation alone is insufficient to account for the experimentally observed loss of selectivity.

In contrast, incorporating an exponential spike-initiation non-linearity produces a modest broadening of tuning curves (Fig. S3C). This effect is substantially amplified when adaptation is combined with non-linear spike initiation, as in the adaptive exponential integrate-and-fire (AdExp) model (Fig. S3D), which reproduces the pronounced broadening observed in food-restricted neurons well.

This behavior can be understood by phase-plane analysis. The exponential term introduces a non-linear voltage nullcline, reshaping the local dynamical landscape near threshold. In this regime, the separation between the voltage (*V*) and adaptation (*w*) nullclines provides a measure of dynamical robustness (Hilgert *et al*., 2025): larger separations imply greater resistance to perturbations from membrane noise, synaptic fluctuations, or parameter variability. Control neurons exhibit a larger nullcline separation, conferring greater robustness and preserving selectivity, whereas food-restricted parameter configurations reduce this separation, rendering the system more susceptible to noise-driven transitions and thereby amplifying tuning curve broadening.

**Figure S3:**
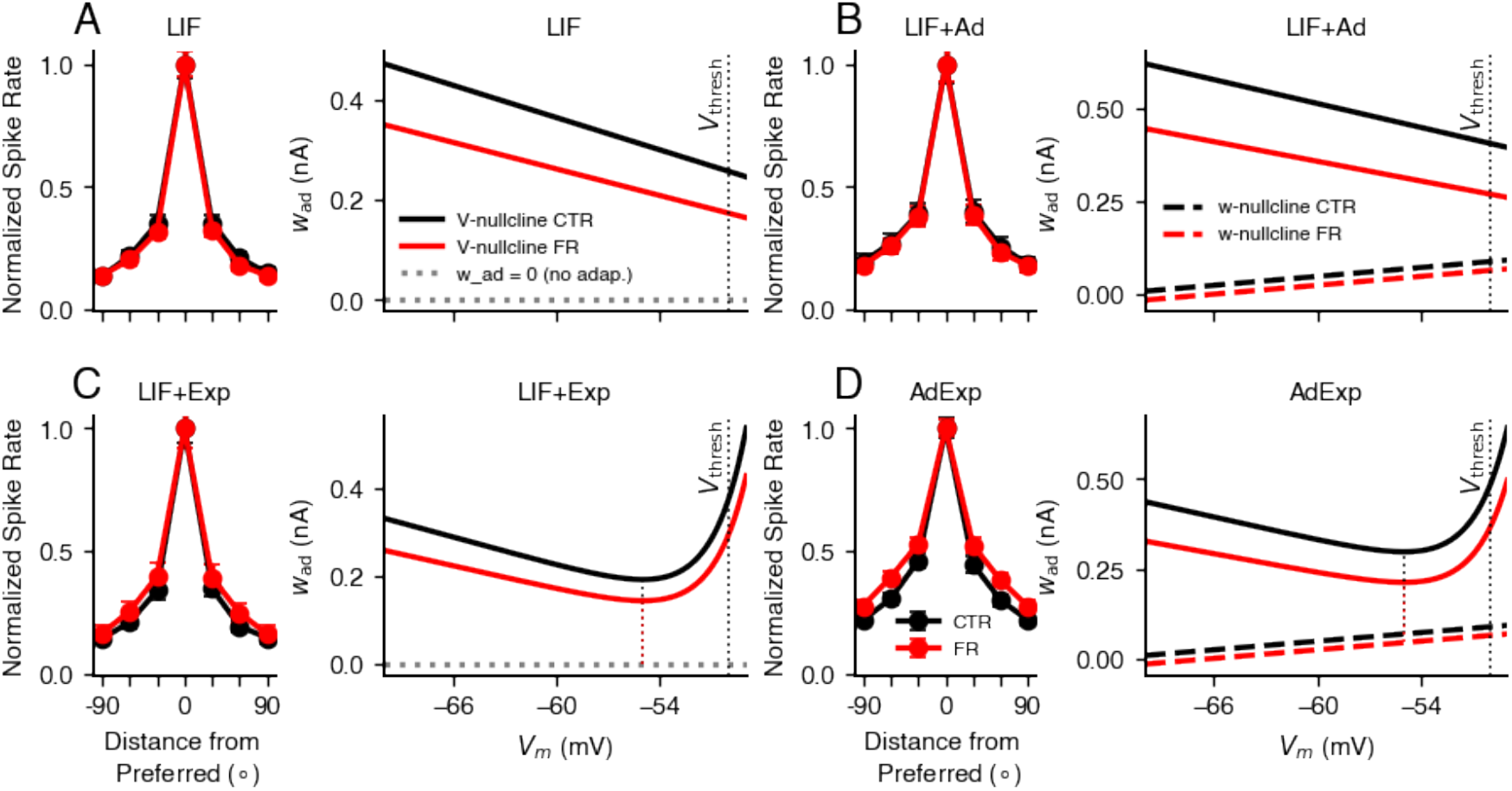
Spike-generation non-linearities and adaptation amplify tuning curve broadening. **A**, A leaky integrate-and-fire (LIF) model with an orientation selectivity task (Fig. S1) does not exhibit tuning curve broadening at experimentally measured membrane noise levels. The corresponding phase plane contains only a linear voltage (*V*) nullcline. **B**, Adding a linear adaptation current to the LIF model does not induce tuning curve broadening; both the *V* - and *w*-nullclines remain linear, **C**, Introducing an exponential spike-initiation term produces a modest broadening of tuning curves. The phase plane now contains a non-linear *V* -nullcline, increasing sensitivity to fluctuations near threshold. **D**, The adaptive exponential integrate-and-fire (AdExp) model reproduces robust tuning curve broadening under experimental noise levels. The combination of a non-linear *V* -nullcline and a linear *w*-nullcline reduces the nullcline separation in food-restricted (FR) neurons relative to control (CTR) neurons, indicating decreased robustness to perturbations and explaining the stronger loss of selectivity in the FR condition.

**Figure S4:**
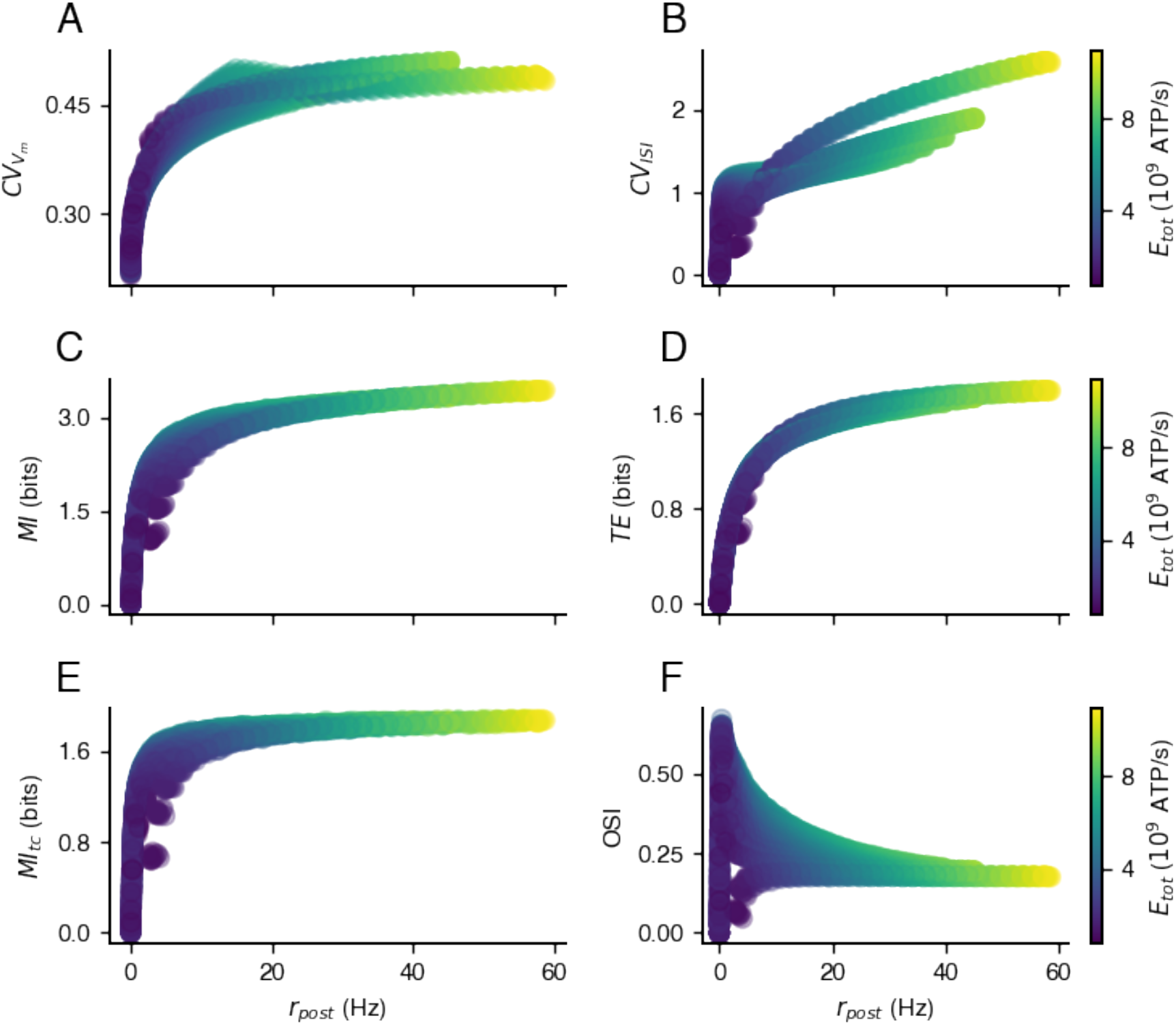
Correlation between firing rate and information measures. Correlation between mean firing rate and **A** coefficient of variation of subthreshold membrane voltage, **B** coefficient of variation of inter spike intervals, **C** mutual information, **D** transfer entropy, and **E** the mutual information based on the tuning curve only for AdExp model simulations with an orientation selectivity task (Fig. S1). **A** – **E** showing a similar relationship as previously reported by (Harris et *al*.,2015). **F**, Orientation selectivity index increases, has a peak at small firing rates and slowly decays as firing rates increase.

## Optimal binsize to maximize MI and TE

**Figure S5:**
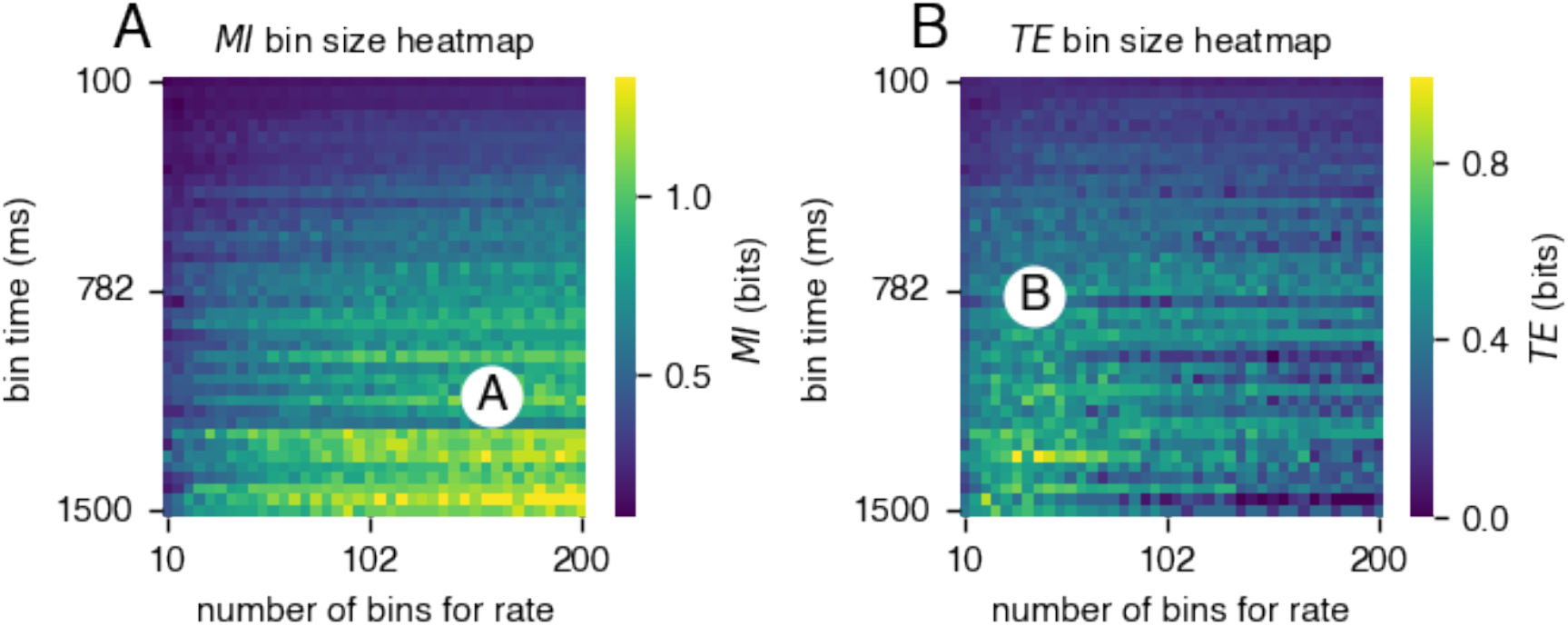
Optimization of the binsize for to calculate mutual information (MI) and transfer entropy (TE) in orientation selectivity tasks (Fig. S1) **A**, Different bin size combinations in time and frequency space were evaluated for mutual information calculation. The white dot marks the optimal discretization parameters: bin time_*MI*_ = 1200 ms and bins rate_*MI*_ = 150. **B**, The same procedure was applied for transfer entropy calculation, yielding optimal parameters of bin time_*TE*_ = 782 ms and bins rate_*TE*_ = 39, with the time lag of *τ* = 294 ms.

## Derivation and rationale behind the energy budget calculation

### Calculating proportionality factor A_**syn**_

To refine the estimate of synaptic transmission energy costs, we computed the contributions of different ionic currents using a detailed compartmental model (Goetz *et al*., 2021) presented in Fig. S6. While previous models assumed that excitatory synaptic currents were dominated by sodium influx, our analysis revealed that calcium currents contribute significantly to the overall energy demand. By quantifying the ratio of sodium to calcium currents and incorporating their distinct ATP requirements for extrusion, we derived a correction factor, *A*_*syn*_, to adjust the energy estimate. Across different stimulus conditions, *A*_*syn*_ remained relatively stable, ranging from 1.055 for non-preferred stimuli to 1.08 for preferred stimuli. To account for inhibitory synaptic energy demands, we set *A*_*syn*_ ≈ 1.1, providing a more accurate estimate of the metabolic cost associated with synaptic transmission.

**Figure S6:**
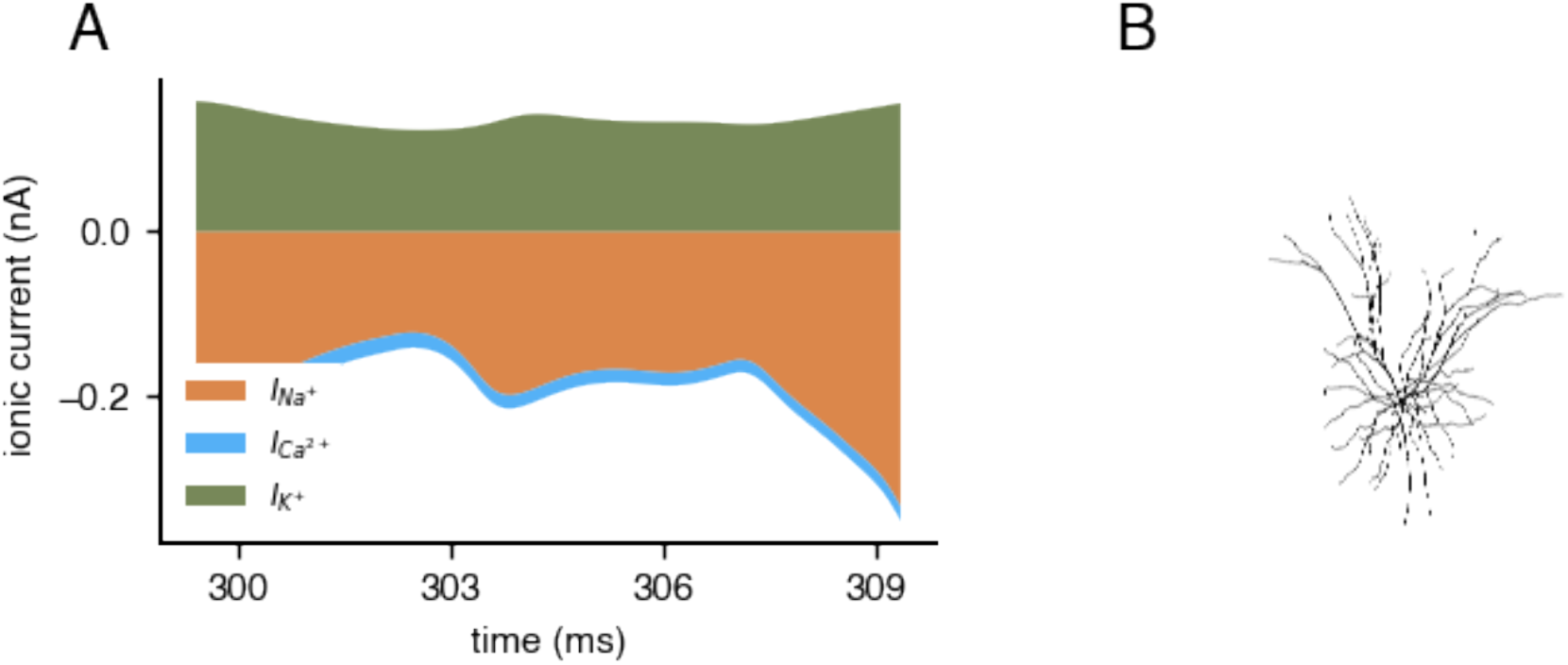
Subthreshold ionic currents of the detailed compartmental model (Goetz *et al*., 2021) **A,** Total synaptic ionic currents summed across all compartments of the detailed model (Goetz *et al*., 2021) over a 10 ms interval. The dominant contributors are the sodium current (*I*_Na_) and potassium current (*I*_K_), while the calcium current (*I*_Ca_) provides a non-negligible contribution to the synaptic energy budget. **B**, Morphology of the detailed compartmental model (Goetz *et al*., 2021).

**Figure S7:**
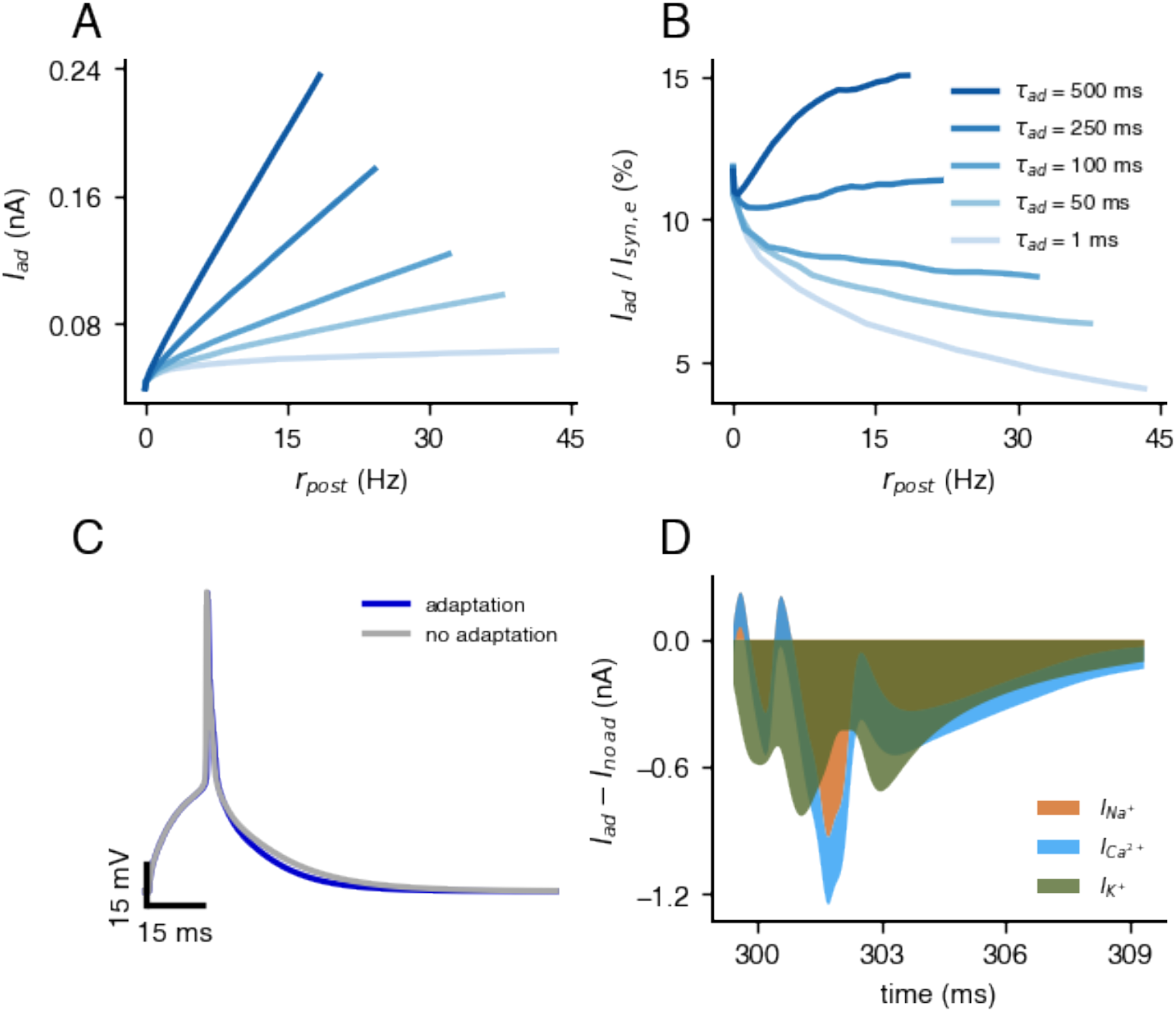
Estimating the cost of adaptation currents. **A**, Mean adaptation current of the AdExp model with given parameters (Tab. 1) as a function of firing rate for four adaptation time constants (1, 50, 100, 500 ms). Higher adaptation time constants lead to larger mean adaptation currents across all firing rates. **B**, Ratio of mean adaptation current to mean excitatory synaptic current as a function of firing rate. For short adaptation time constants (1–100 ms), this ratio decreases with firing rate, while for a long time constant (500 ms), the ratio increases (indicating greater metabolic cost at higher activity levels). **C**, Membrane voltage traces from the detailed compartmental model (Goetz *et al*., 2021) in response to a brief step current (100 ms onset, 15 ms duration). Simulations with and without adaptation currents (*I*_*M*_ and 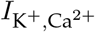) are compared. Without adaptation, the post-spike depolarization is elevated. **D**, Ionic currents 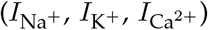 recorded following spike initiation (from 1.5 ms post-spike). Disabling adaptation currents increases net 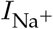 and 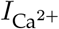 (also 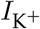) influx, thereby raising the energetic cost per spike.

### Derivation of resting potential energy demand

In the following we perform the derivation of formula formula (4) in Attwell and Laughlin (2001) for the consumed energy *E*_*RP*_ (ATP per second) of a neuron as a function of the resting membrane potential *V*_*RP*_, the reverse potentials *V*_Na_ and *V*_K_, the Faraday constant *F*, and the input resistance *R*_*m*_. The given relationships are:

I. Ionic currents at rest: *I*_*pump*_ = *g*_Na_(*V*_Na_ − *V*_*RP*_)+ *g*_K_(*V*_K_ − *V*_*RP*_)
II. Total consumption of ATP/time by Na^+^/K^+^-pump: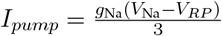
III. 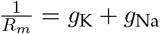
IV. 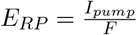

The steps of derivation are:

#### Step 1: Set the equations equal to each other

Equate the two expressions for *I*_*pump*_:

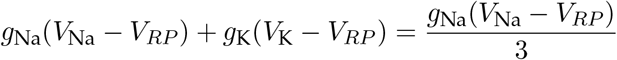

#### Step 2: Simplify the equation

Multiply both sides by 3 to eliminate the fraction:

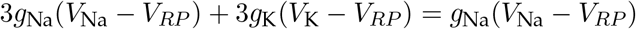

Subtract *g*_Na_(*V*_Na_ − *V*_*RP*_) from both sides:

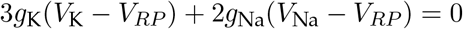

#### Step 3: Rearrange to isolate *V*_*RP*_

Distribute the terms:

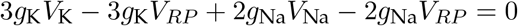

Combine like terms:

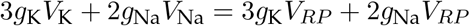

Factor out *V*_*RP*_ on the right side:

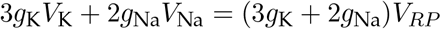

Solve for *V*_*RP*_ :

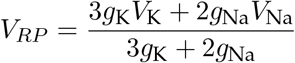

Thus, the resting membrane potential *V*_*RP*_ is given by this equation.

#### Step 4: Rearrange to isolate *g*_**Na**_ **and** *g*_**K**_

Distribute the terms:

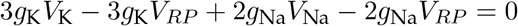

Combine like terms:

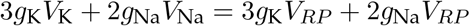

Rearrange to separate *g*_Na_ and *g*_K_:

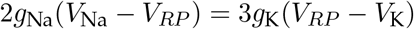

Solve for *g*_Na_:

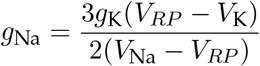

#### Step 5: Use the input resistance relationship to solve for *g*_K_ (III)

Using the relationship 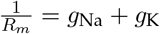:

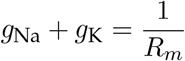

Substitute the expression for *g*_Na_:

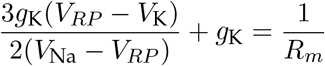

Factor out *g*_K_:

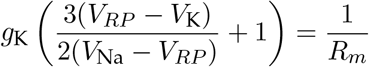

Combine terms inside the parentheses:

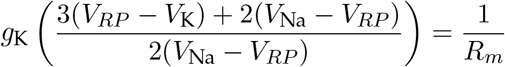

Simplify the numerator:

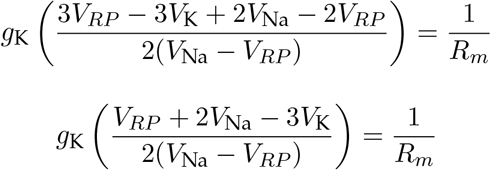

Solve for *g*_K_:

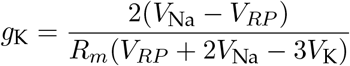

#### Step 6: Substitute *g*_**K**_ **back into the expression for** *g*_**Na**_

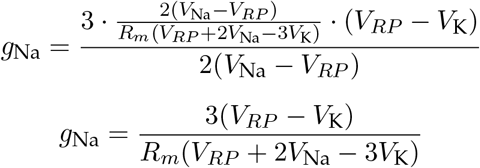

#### Step 7: Substitute *I*_*pump*_ into *E*_*RP*_ (II in IV)

From the given relationships:

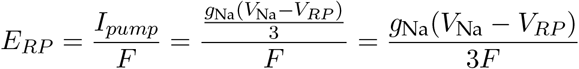

#### Step 8: Substitute *g*_**Na**_ **in the energy consumption equation** *E*_*RP*_

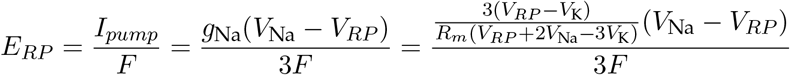

Simplify:

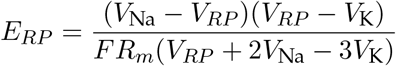

and arrive at the final formula (4) given by (Attwell and Laughlin, 2001). Alternative formulation for better understanding of the derivation:

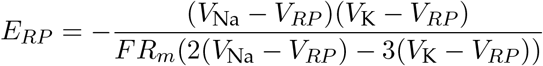

### Estimating the cost of the resting potential with additional ionic conductances

Here we provide an informative derivation of the effect of including an inward-hyperpolarisin (*I*_*h*_) conductance, mediated by hyperpolarization-activated cyclic nucleotide–gated (HCN) channels, on the ATP cost of maintaining a given membrane resting potential. As neuronal membranes are maintained by a large number of conductances (Goaillard and Marder, 2021), this derivation could be extended further. The above assumptions are modified as followed:

1. Ionic currents at rest. The pump currents now consist of an additional term to account for the *I*_*h*_, so that:

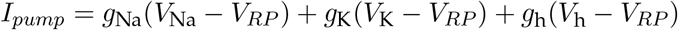

The reversal potential of the *I*_*h*_ current, *V*_h_, is taken to be −43mV (McCormick and Pape, 1990).
2. As the *I*_*h*_ current is a mixed cation current typically assumed to consist of ∼ 3*/*4 Na^+^ (McCormick and Pape, 1990), the total consumption of ATP/time by the Na^+^/K^+^-pump is:

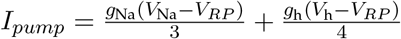
3. The membrane resistance now depends on the sum of the three conductances:

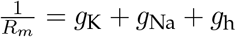
4. As before:

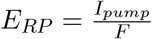

Now, the above system is underconstrained in terms of ionic conductances. We introduce a parameter *α* to describe the ratio of *g*_Na_ to *g*_K_ so that *g*_Na_ = *αg*_K_. Typical estimates of *α* are in the region of 0.05 (Chrysafides S M and S, 2024). This relationship could change with the resting potential, but the Na^+^ conductance will typically remain far lower than the K^+^ conductance at rest. It is then possible to use the first two relationships above to write *g*_K_ in terms of *g*_h_ for a given resting potential:

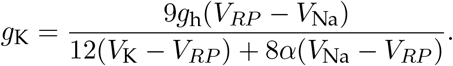

The membrane resistance relationship can then be used to write the three ionic conductances

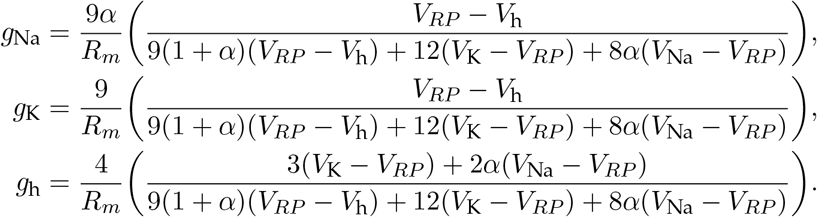

This results in the energy to maintain the resting potential:

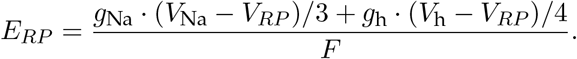

These conductances can be plotted individually (Fig. S8A) and show how, in contrast to the model with only two conductances, it is possible for sodium channels to close as the resting potential increases. This further implies that both increasing the membrane resistance *R*_*m*_ and the resting potential *V*_RP_ can save energy (Fig. S8B). The joint effect of the experimentally observed changes could therefore lead to a more substantial ATP saving given the extra degree of freedom provided by the HCN channels (Fig. S8C). In reality, many other channels are involved in maintaining the resting potential and we hypothesise that energy savings here could be even greater. Clarifying the exact mechanisms involved in these changes would require further experiments.

**Figure S8:**
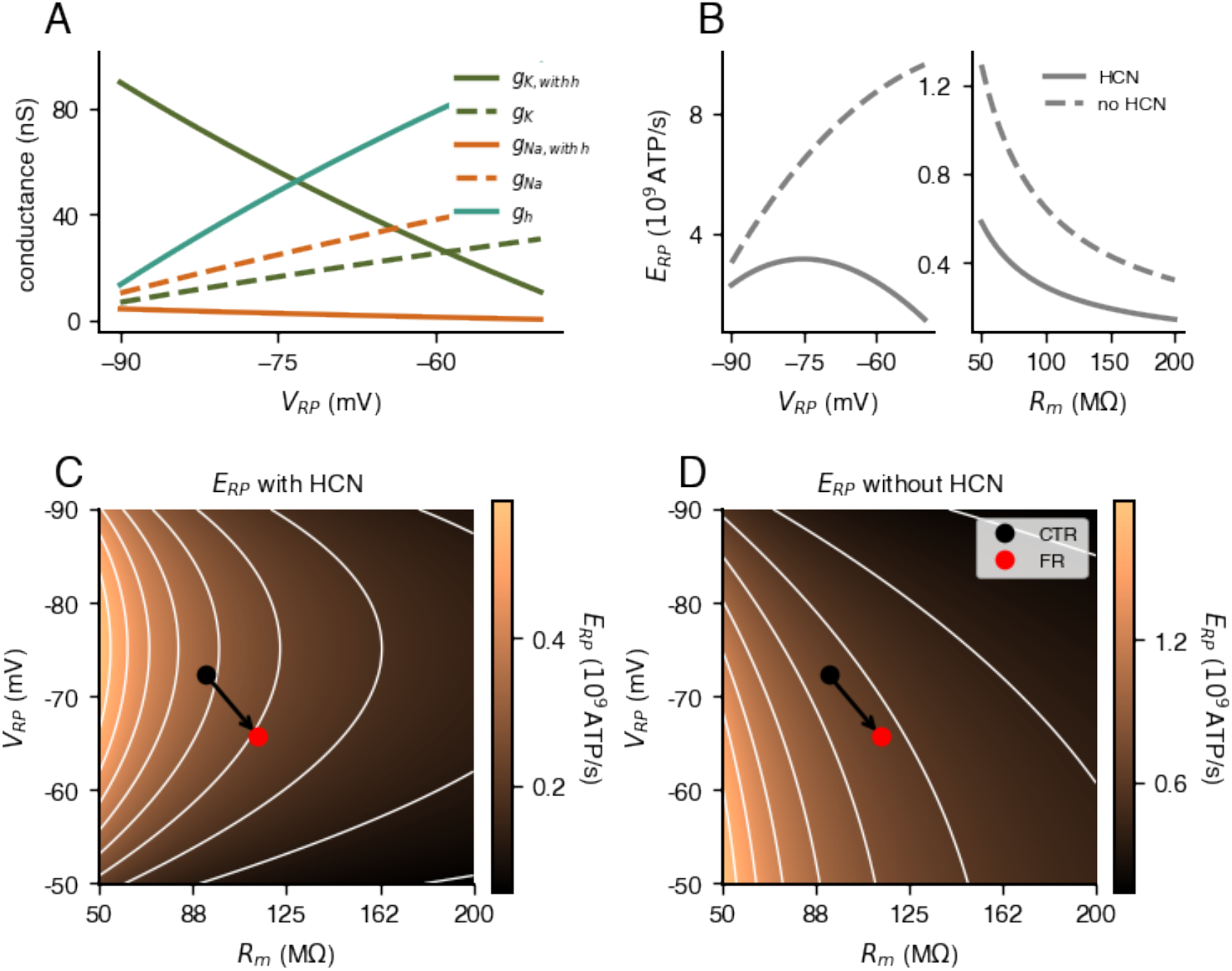
Effect of considering additional ionic conductances of estimates of ATP consumption. **A**, Individual ionic conductances as a function of resting potential with (solid lines) and without (dashed lines) considering *I*_*h*_. Input resistance is 92.3M Ω. **B**, ATP consumption necessary to maintain membrane properties as a function of resting potential (left, *R*_*m*_ = 92.3MΩ) and input resistance (right, *V*_*RP*_ = − 72.3mV) with (solid lines) and without (dashed lines) considering *I*_*h*_. **C**, ATP consumption necessary to maintain membrane properties as a function of both resting potential and input resistance with (left) and without (right) considering *I*_*h*_. Mean values for the control (black) and food restricted (red) cells are highlighted.

**Figure S9:**
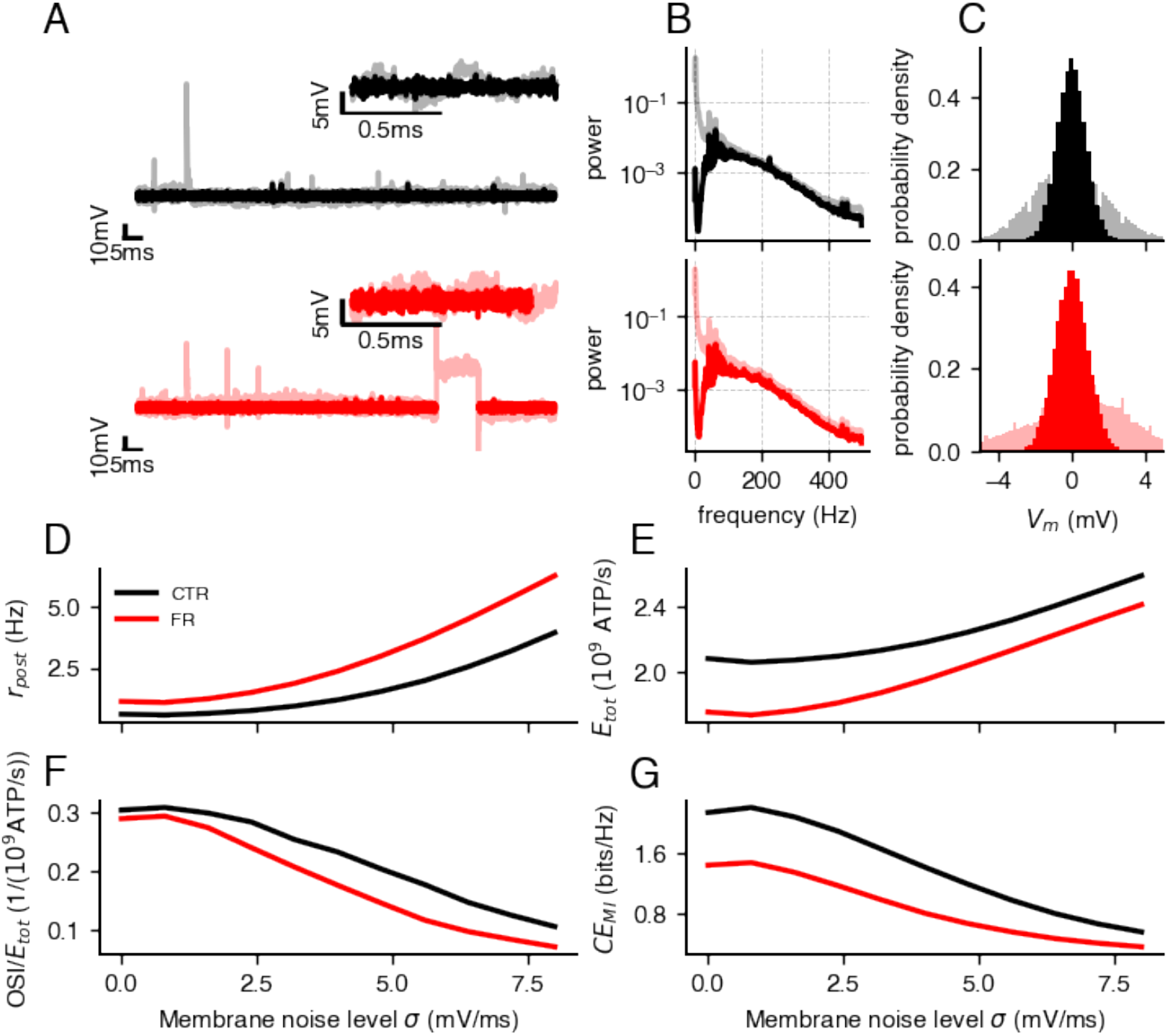
Experimental membrane noise amplitude extraction and effects of different membrane noise levels on neuronal behavior of cells from control (CTR) and food-restricted (FR) animals. **A**, Preprocessing pipeline for membrane noise analysis: large deviations are excluded by windowing, followed by Notch-filtering to remove low-frequency oscillations, and normalization using a moving average to correct for slow baseline drifts. **B**, Power spectral density of membrane potential fluctuations, illustrating the frequency components of the signal. **C**, Probability density distribution of membrane potential fluctuations, used to estimate the standard deviation of intrinsic noise. **D**, Mean firing rate rises exponentially with increasing membrane noise for simulations of AdExp models with an orientation selectivity task (Fig. S1). **E**, Total energy expenditure mirrors the increase of firing rate but CTR energy demand is constantly higher than FR. **F**, Orientation selectivity per energy as a function of membrane noise, including non-spiking trials. **G**, Coding efficiency (mutual information per spike) mirrors the orientation selectivity index (OSI), showing an initial increase at low membrane noise levels due to stochastic resonance, followed by a monotonic decline at higher noise.

**Figure S10:**
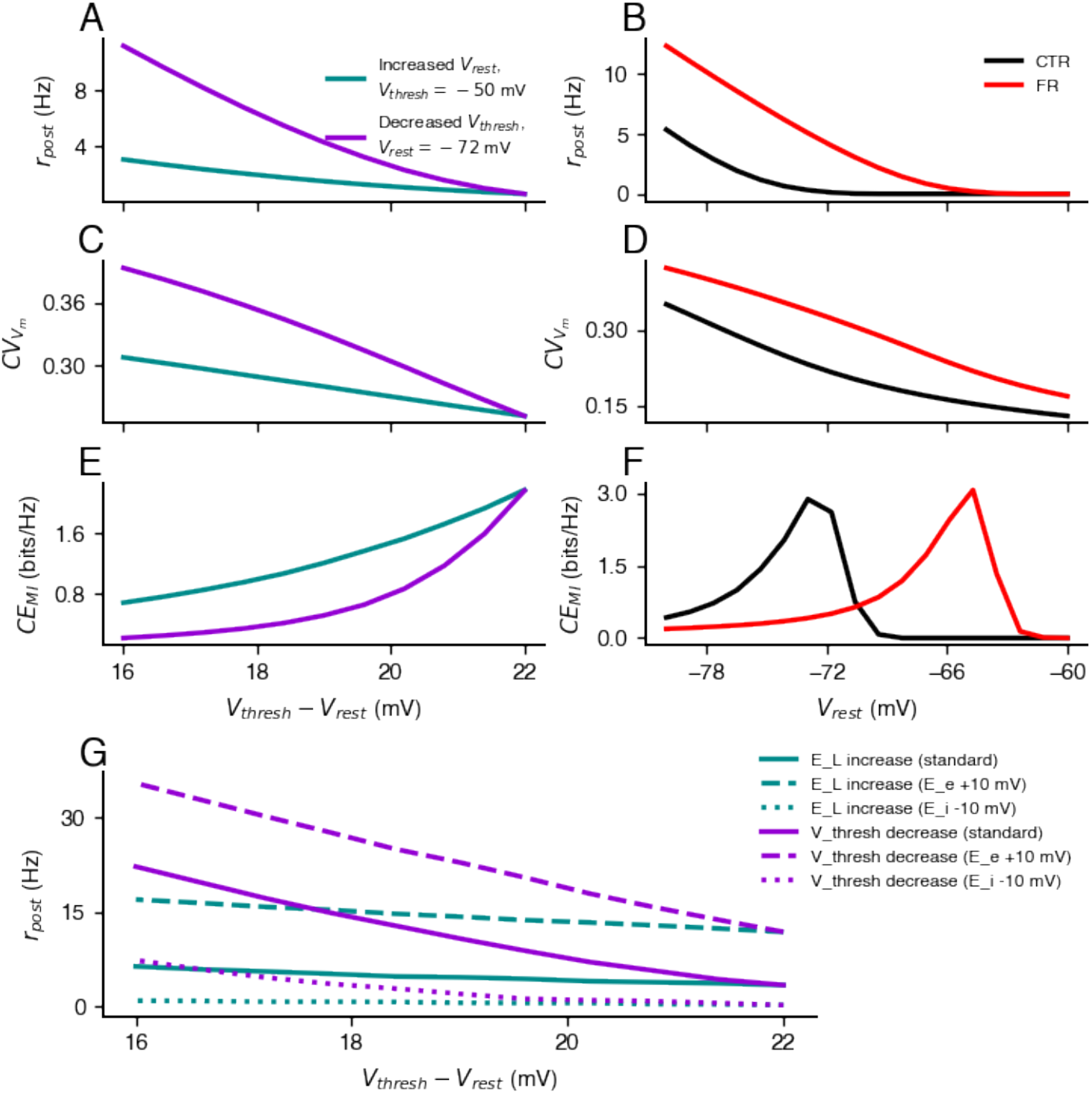
Resting potential and synaptic reversal values shape firing rate, energy use, and information transmission (AdExp model with an orientation selectivity task (Fig. S1)). **A**, Postsynaptic firing rate, **B**, mutual information, **C**, and mutual information per spike (coding efficiency) as functions of the voltage gap between threshold and resting potential, modulated either by depolarizing the resting potential (turquoise) or lowering the threshold (purple). Lower thresholds produce higher firing rates and, consequently, higher mutual information, though at increased energetic cost. Energy consumption scales approximately linearly with firing rate at high rates. But the coding-efficiency (mutual information per spike) is higher for a decreased resting potential. **D-F**, Sliding a fixed resting–threshold gap across the membrane potential range reveals that food-restricted cells (red) maintain **D**, higher firing rates, **E**, higher information content. However, at more depolarized resting potentials (*>* –70 mV), control cells become more energy-efficient. **F**, Similarly to the higher energy efficiency, the coding-efficiency is higher for low resting potentials in control cells and higher for depolarized resting potentials in food-restricted cells.

**Figure S11:**
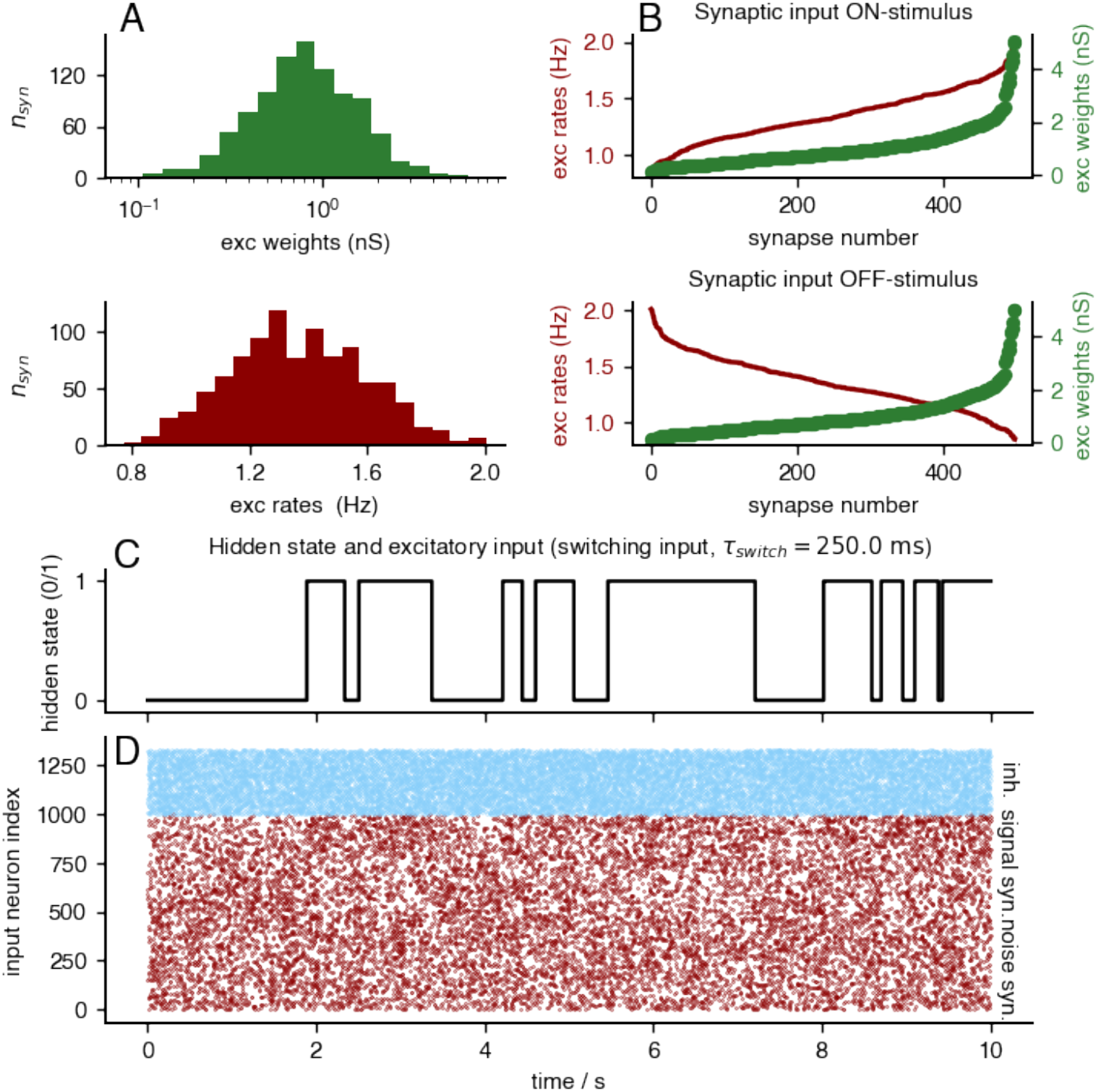
Design of Markov-process switching ON-OFF stimulus excitatory and inhibitory synaptic input similar to the stimulation protocol of Zeldenrust *et al*. (2024). **A – Top**, Excitatory synaptic weights follow a log-normal distribution 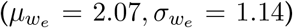 **B– Bottom**, Excitatory presynaptic firing rates are normally distributed 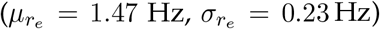. **B**, The preferred ON and non-preferred OFF stimuli assigned synaptic inputs based on the matching scheme, visualized as cityscape histograms: presynaptic firing rates (dark red) as a function of synaptic strength (green). The preferred stimulus (ON) elicits the strongest total synaptic drive, whereas non-preferred stimulus (OFF) exhibit the weakest. **C**, Hidden state representing the ON-OFF stimulus switching task. **D**, Raster plot of synaptic input times, distinguishing inhibitory (light blue excitatory signal (dark red), and excitatory noise (dark red).

**Figure S12:**
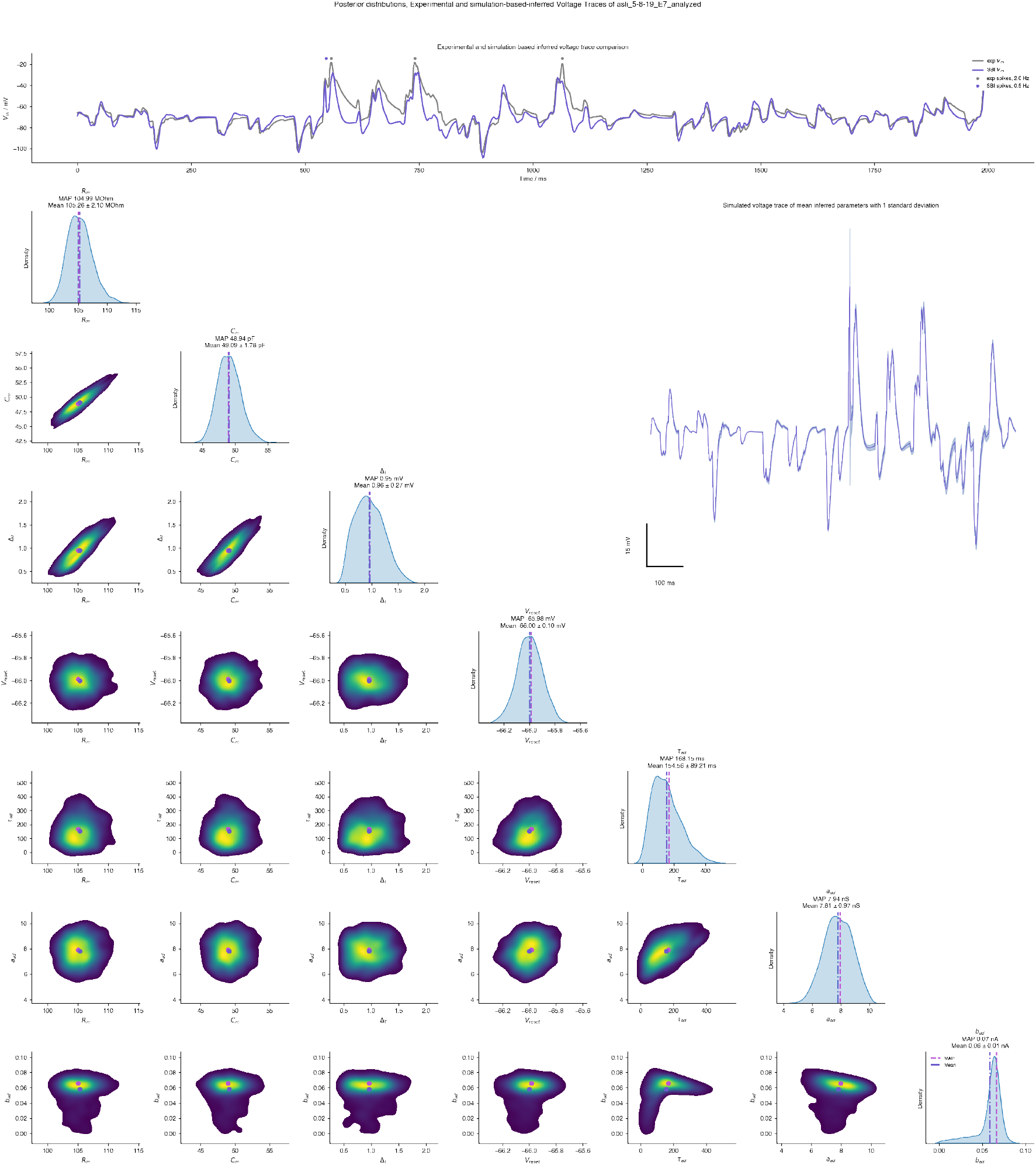
SBI posterior distributions and validation of inferred parameters. **Top row,** Experimental voltage trace (Zeldenrust *et al*., 2024) compared to a simulated trace generated using the posterior mean parameter estimates. **Right**, Voltage traces simulated using parameters sampled at the posterior mean ± 1 standard deviation, illustrating uncertainty in the inferred dynamics, **Left**, One- and two-dimensional marginal posterior densities for inferred parameters, with posterior means (blue) and maximum a posteriori (MAP — purple) estimates indicated.

**Figure S13:**
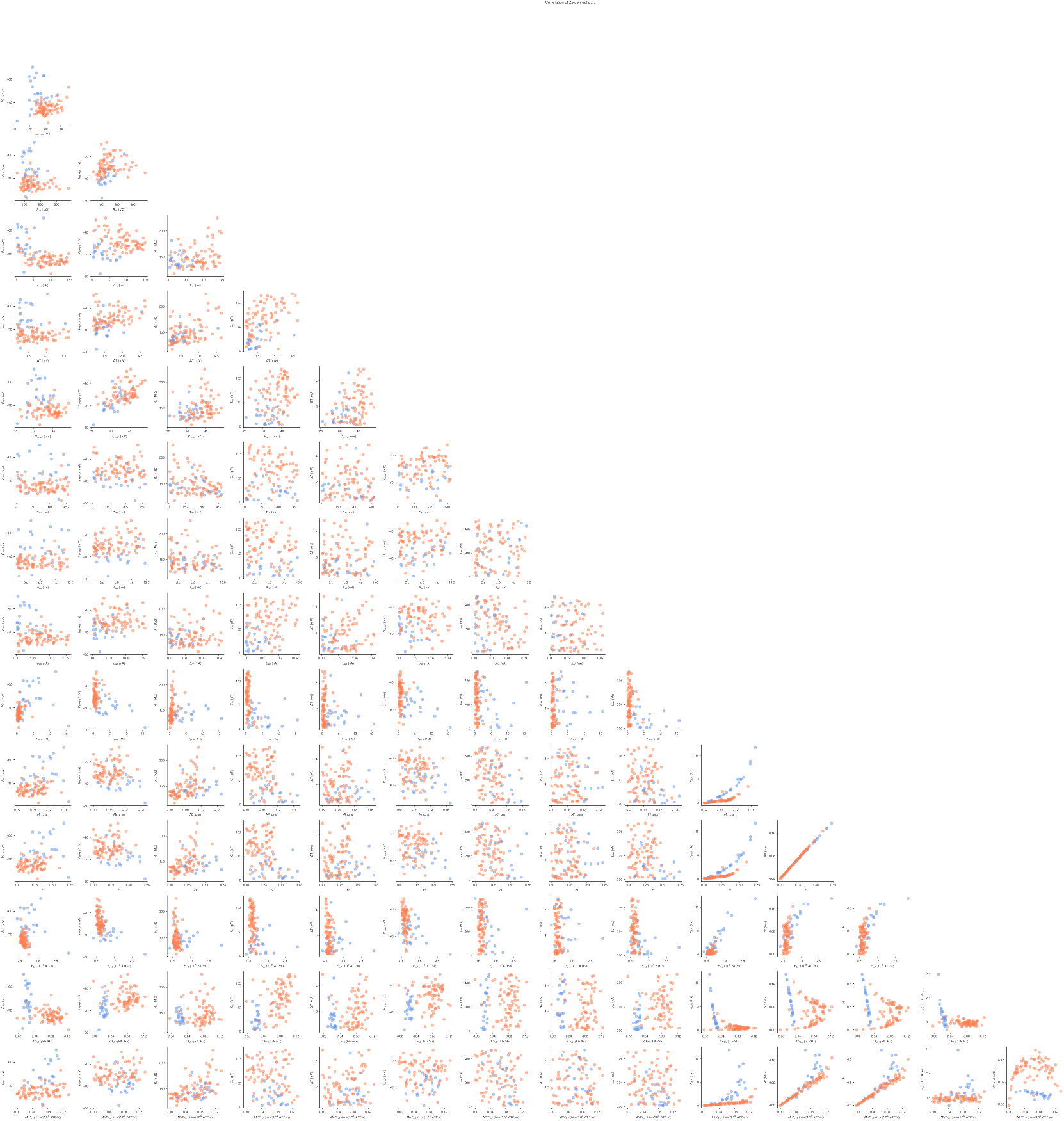
Correlation structure of excitatory and inhibitory cell parameters and functional properties. Pairwise correlations among electrophysiological parameters and derived properties such as mutual information and total energy consumption from Zeldenrust *et al*. (2024) are shown separately for excitatory and inhibitory neurons.

**Figure S14:**
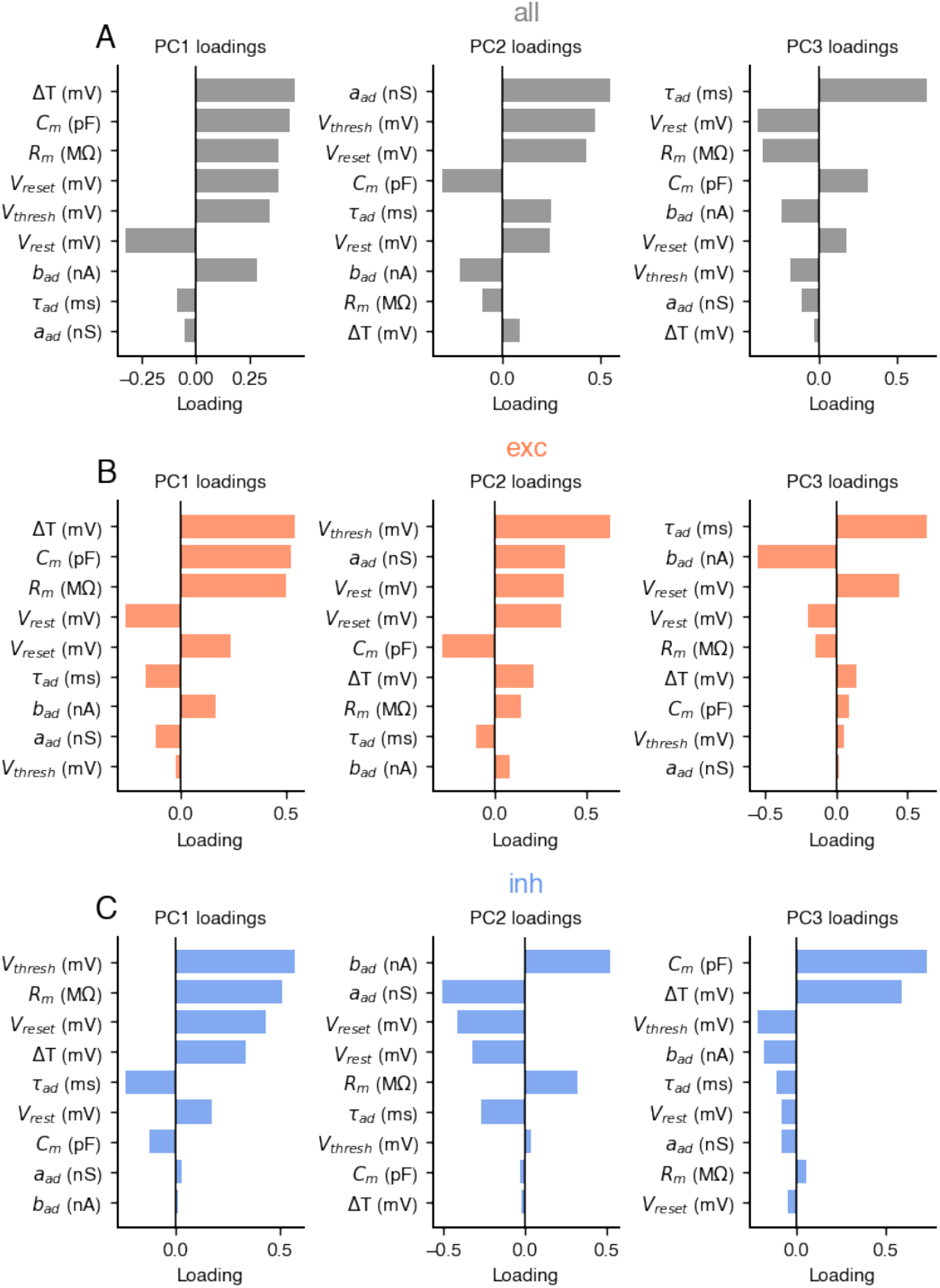
Principal component loadings (PC1–3) for all, excitatory, and inhibitory neurons. **A**, Loadings of the first three principal components (PC1–3) across all neurons, capturing the dominant axes of electrophysiological variability in our parameter inference of experimental data from Zelden-rust et al. (2024). **B**, PC1–3 loadings restricted to excitatory neurons, highlighting excitatory-specific variance structure. **C**, PC1–3 loadings restricted to inhibitory neurons, revealing distinct variance contributions relative to excitatory cells.

## Notes

https://github.com/trieschlab/energy-information-trade-off

### Competing Interest Statement

The authors have declared no competing interest.

### Summary of Updates

Abstract and title updated to clarify, Supplemental figures updated.

## References

Alle H, Roth A, Geiger JRP (2009) Energy-Efficient Action Potentials in Hippocampal Mossy Fibers. Science 325:1405–1408.

Altschuler SJ, Wu LF (2010) Cellular Heterogeneity: Do Differences Make a Difference? Cell 141:559–563.

Attwell D, Laughlin SB (2001) An Energy Budget for Signaling in the Grey Matter of the Brain. Journal of Cerebral Blood Flow & Metabolism 21:1133–1145.

Balasubramanian V (2021) Brain power. Proceedings of the National Academy of Sciences 118:e2107022118.

Barlow HB (1961) Possible principles underlying the transformations of sensory messages. Sensory communication 1:217–233.

Barral J, Reyes AD (2016) Synaptic scaling rule preserves excitatory–inhibitory balance and salient neuronal network dynamics. Nature Neuroscience 19:1690–1696.

Bazzigaluppi P, Amini AE, Weisspapir I, Stefanovic B, Carlen P (2017) Hungry Neurons: Metabolic Insights on Seizure Dynamics. International Journal of Molecular Sciences 18:2269.

Beaulieu-Laroche L, Brown NJ, Hansen M, Toloza EHS, Sharma J, Williams ZM, Frosch MP, Cosgrove GR, Cash SS, Harnett MT (2021) Allometric rules for mammalian cortical layer 5 neuron biophysics. Nature 600:274–278.

Bird AD, Richardson MJ (2018) Transmission of temporally correlated spike trains through synapses with short-term depression. PLOS Computational Biology 14:e1006232.

Braitenberg V, Schüz A (1998) Cortex: statistics and geometry of neuronal connectivity Springer Science & Business Media.

Brette R, Gerstner W (2005) Adaptive Exponential Integrate-and-Fire Model as an Effective Description of Neuronal Activity. Journal of Neurophysiology 94:3637–3642.

Brunel N, van Rossum MC (2007) Lapicque’s 1907 paper: from frogs to integrate-and-fire. Biological cybernetics 97:337–339.

Buzsaki G, Kaila K, Raichle M (2007) Inhibition and Brain Work. Neuron 56:771–783.

Buzsaki G, Mizuseki K (2014) The log-dynamic brain: how skewed distributions affect network operations. Nature Reviews Neuroscience 15:264–278.

Carter BC, Bean BP (2009) Sodium Entry during Action Potentials of Mammalian Neurons: Incomplete Inactivation and Reduced Metabolic Efficiency in Fast-Spiking Neurons. Neuron 64:898–909.

Chen BL, Hall DH, Chklovskii DB (2006) Wiring optimization can relate neuronal structure and function. Proceedings of the National Academy of Sciences 103:4723–4728.

Chrysafides S M B SJ, S S (2024) Physiology, resting potential. StatPearls.

Dahmen D, Hutt A, Indiveri G, Kennedy A, Lefebvre J, Mazzucato L, Motter AE, Narayanan R, Payvand M, Planert H, Gast R (2025) How heterogeneity shapes dynamics and computation in the brain. Neuron.

Deistler M, Macke JH, Gonçalves PJ (2022) Energy-efficient network activity from disparate circuit parameters. Proceedings of the National Academy of Sciences 119:e2207632119.

Del Castillo J, Katz B (1954) Statistical factors involved in neuromuscular facilitation and depression. The Journal of Physiology 124:574.

Deneve S, Machens CK (2016) Efficient codes and balanced networks. Nature Neuroscience 19:375–382.

Desai NS, Rutherford LC, Turrigiano GG (1999) Plasticity in the intrinsic excitability of cortical pyramidal neurons. Nature Neuroscience 2:515–520.

Dimakou A, Pezzulo G, Zangrossi A, Corbetta M (2025) The predictive nature of spontaneous brain activity across scales and species. Neuron 113:1310–1332.

Dougherty KA, Islam T, Johnston D (2012) Intrinsic excitability of CA1 pyramidal neurones from the rat dorsal and ventral hippocampus. The Journal of Physiology 590:5707–5722.

Duarte R, Morrison A (2019) Leveraging heterogeneity for neural computation with fading memory in layer 2/3 cortical microcircuits. PLoS Computational Biology 15:e1006781.

Edelman GM, Gally JA (2001) Degeneracy and complexity in biological systems. Proceedings of the National Academy of Sciences of the United States of America 98:13763–8.

Engl E, Attwell D (2015) Non signalling energy use in the brain. The Journal of Physiology 593:3417–3429.

Gast R, Solla SA, Kennedy A (2024) Neural heterogeneity controls computations in spiking neural networks. Proceedings of the National Academy of Sciences 121:e2311885121.

Gauld OM, Packer AM, Russell LE, Dalgleish HW, Iuga M, Sacadura F, Roth A, Clark BA, Häusser M (2024) A latent pool of neurons silenced by sensory-evoked inhibition can be recruited to enhance perception. Neuron 112:2386–2403.e6.

Gerstner W, Kistler WM, Naud R, Paninski L (2014) Neuronal Dynamics: From single neurons to networks and models of cognition Cambridge University Press, Cambridge.

Giudice MD, Crespi BJ (2018) Basic functional trade-offs in cognition: An integrative framework. Cognition 179:56–70.

Goaillard JM, Marder E (2021) Ion Channel Degeneracy, Variability, and Covariation in Neuron and Circuit Resilience. Annual Review of Neuroscience 44:1–23.

Goetz L, Roth A, Hausser M (2021) Active dendrites enable strong but sparse inputs to determine orientation selectivity. Proceedings of the National Academy of Sciences 118:e2017339118.

Goldman M (2004) Enhancement of information transmission efficiency by synaptic failures. Neural Computation 16:1137–1162.

Greenberg DS, Nonnenmacher M, Macke JH (2019) Automatic posterior transformation for likelihood-free inference. Proceedings of the 36th International Conference on Machine Learning (ICML) pp. 2404–2414.

Haggard M, Chacron MJ (2023) Coding of object location by heterogeneous neural populations with spatially dependent correlations in weakly electric fish. PLOS Computational Biology 19:e1010938.

Haider B, Häusser M, Carandini M (2013) Inhibition dominates sensory responses in the awake cortex. Nature 493:97–100.

Harris JJ, Jolivet R, Engl E, Attwell D (2015) Energy-Efficient Information Transfer by Visual Pathway Synapses. Current Biology 25:3151–3160.

Harris JJ, Engl E, Attwell D, Jolivet RB (2019) Energy-efficient information transfer at thalamocortical synapses. PLoS Computational Biology 15:e1007226.

Harris J, Jolivet R, Attwell D (2012) Synaptic Energy Use and Supply. Neuron 75:762–777.

Hilgert S, Hannah S, Niemeyer N, Huthmacher L, Hürkey S, Schleimer J, Schreiber S, Duch C, Ryglewski S (2025) Robustness through variability: ion channel isoform diversity safeguards neuronal excitability. bioRxiv p. 2025.09.16.676508.

Hodgkin AL, Huxley AF (1952) A quantitative description of membrane current and its application to conduction and excitation in nerve. The Journal of Physiology 117:500–544.

Hodgkin JA (1975) The optimum density of sodium channels in an unmyelinated nerve. Philosophical Transactions of the Royal Society of London. B, Biological Sciences 270:297–300.

Howarth C, Gleeson P, Attwell D (2012) Updated Energy Budgets for Neural Computation in the Neocortex and Cerebellum. Journal of Cerebral Blood Flow & Metabolism 32:1222–1232.

Hyder F, Rothman DL, Bennett MR (2013) Cortical energy demands of signaling and nonsignaling components in brain are conserved across mammalian species and activity levels. Proceedings of the National Academy of Sciences 110:3549–3554.

Indiveri G (2025) Neuromorphic is dead. long live neuromorphic. Neuron 113:3311–3314.

Isler K, Schaik CPv (2009) The Expensive Brain: A framework for explaining evolutionary changes in brain size. Journal of Human Evolution 57:392–400.

Jedlicka P, Bird AD, Cuntz H (2022) Pareto optimality, economy–effectiveness trade-offs and ion channel degeneracy: improving population modelling for single neurons. Open Biology 12:220073.

Joshi N, Burg SvD, Celikel T, Zeldenrust F (2025) Neuronal identity is not static: An input-driven perspective. PLOS Computational Biology 21:e1013821.

Klemmensen MM, Borrowman SH, Pearce C, Pyles B, Chandra B (2024) Mitochondrial dysfunction in neurodegenerative disorders. Neurotherapeutics 21:e00292.

Kumari S, Narayanan R (2026) Intrinsic excitability of rat hippocampal granule cells increases along the dorsal-to-ventral axis. bioRxiv p. 2026.01.01.697271.

Kohler-Solis A, Schirmeier S (2025) Neuronal metabolism: Surprisingly flexible. Current Biology 35:R1098–R1101.

Laughlin SB, Sejnowski TJ (2003) Communication in Neuronal Networks. Science 301:1870–1874.

Le Masson G, Przedborski S, Abbott L (2014) A Computational Model of Motor Neuron Degeneration. Neuron 83:975–988.

Lennie P (2003) The Cost of Cortical Computation. Current Biology 13:493–497.

Levy WB, Baxter RA (1996) Energy Efficient Neural Codes. Neural Computation 8:531–543.

Levy WB, Baxter RA (2002) Energy-Efficient Neuronal Computation via Quantal Synaptic Failures. The Journal of Neuroscience 22:4746–4755.

Lezmy J, Harris JJ, Attwell D (2021) Optimising the energetic cost of the glutamatergic synapse. Neuropharmacology 197:108727.

Li HL, van Rossum MC (2020) Energy efficient synaptic plasticity. eLife 9:e50804.

Li S, Sheng ZH (2022) Energy matters: presynaptic metabolism and the maintenance of synaptic transmission. Nature Reviews Neuroscience 23:4–22.

Lueckmann JM, Boelts J, Greenberg DS, Gonçalves PJ, Macke JH (2021) Benchmarking simulation-based inference. Proceedings of the National Academy of Sciences 118:e2103070118.

Marder E, Goaillard JM (2006) Variability, compensation and homeostasis in neuron and network function. Nature Reviews Neuroscience 7:563–574.

Marder E, Prinz AA (2002) Modeling stability in neuron and network function: the role of activity in homeostasis. BioEssays 24:1145–1154.

McCormick DA, Pape HC (1990) Properties of a hyperpolarization-activated cation current and its role in rhythmic oscillation in thalamic relay neurones. The Journal of Physiology 431:291–318.

Mejias JF, Longtin A (2012) Optimal heterogeneity for coding in spiking neural networks. Physical Review Letters 108:228102.

Mishra P, Narayanan R (2021) Ion-channel degeneracy: Multiple ion channels heterogeneously regulate intrinsic physiology of rat hippocampal granule cells. Physiological Reports 9:e14963.

Navarrete A, Schaik CPv, Isler K (2011) Energetics and the evolution of human brain size. Nature 480:91–93.

Niven JE, Laughlin SB (2008) Energy limitation as a selective pressure on the evolution of sensory systems. Journal of Experimental Biology 211:1792–1804.

N’dri AW, Barbier T, Teuliere C, Triesch J (2025) Predictive coding light. Nature Communications 16:8880.

O’Leary T, Williams AH, Caplan JS, Marder E (2013) Correlations in ion channel expression emerge from homeostatic tuning rules. Proceedings of the National Academy of Sciences 110:E2645–E2654.

Olshausen BA, Field DJ (1996) Emergence of simple-cell receptive field properties by learning a sparse code for natural images. Nature 381:607–609.

Pache A, van Rossum MC (2023) Energetically efficient learning in neuronal networks. Current Opinion in Neurobiology 83:102779.

Padamsey Z, Katsanevaki D, Dupuy N, Rochefort NL (2022) Neocortex saves energy by reducing coding precision during food scarcity. Neuron 110:280–296.e10.

Padamsey Z, Rochefort NL (2023) Paying the brain’s energy bill. Current Opinion in Neurobiology 78:102668.

Padmanabhan K, Urban NN (2010) Intrinsic biophysical diversity decorrelates neuronal firing while increasing information content. Nature Neuroscience 13:1276–1282.

Pallasdies F, Norton P, Schleimer JH, Schreiber S (2021) Neural optimization: Understanding trade-offs with Pareto theory. Current Opinion in Neurobiology 71:84–91.

Perez-Nieves N, Leung VCH, Dragotti PL, Goodman DFM (2021) Neural heterogeneity promotes robust learning. Nature Communications 12:5791.

Prinz AA, Billimoria CP, Marder E (2003) Alternative to Hand-Tuning Conductance-Based Models: Construction and Analysis of Databases of Model Neurons. Journal of Neurophysiology 90:3998–4015.

Prinz AA, Bucher D, Marder E (2004) Similar network activity from disparate circuit parameters. Nature Neuroscience 7:1345–1352.

Raimondo JV, Richards BA, Woodin MA (2017) Neuronal chloride and excitability — the big impact of small changes. Current Opinion in Neurobiology 43:35–42.

Remme MWH, Rinzel J, Schreiber S (2018) Function and energy consumption constrain neuronal biophysics in a canonical computation: Coincidence detection. PLoS Computational Biology 14:e1006612.

Rich S, Chameh HM, Lefebvre J, Valiante TA (2022) Loss of neuronal heterogeneity in epileptogenic human tissue impairs network resilience to sudden changes in synchrony. Cell Reports 39:110863.

Riveros N, Fiedler J, Lagos N, Munoz C, Orrego F (1986) Glutamate in rat brain cortex synaptic vesicles: influence of the vesicle isolation procedure. Brain Research 386:405–408.

Savtchenko LP, Sylantyev S, Rusakov DA (2013) Central synapses release a resource-efficient amount of glutamate. Nature Neuroscience 16:10–12.

Schneider M, Bird AD, Gidon A, Triesch J, Jedlicka P, Cuntz H (2023) Biological complexity facilitates tuning of the neuronal parameter space. PLoS computational biology 19:e1011212.

Schreiber S, Machens CK, Herz AVM, Laughlin SB (2002) Energy-Efficient Coding with Discrete Stochastic Events. Neural Computation 14:1323–1346.

Schreiber T (2000) Measuring Information Transfer. Physical Review Letters 85:461–464.

Sengupta B, Laughlin SB, Niven JE (2013) Balanced Excitatory and Inhibitory Synaptic Currents Promote Efficient Coding and Metabolic Efficiency. PLoS Computational Biology 9:e1003263.

Sengupta B, Stemmler M, Laughlin SB, Niven JE (2010) Action Potential Energy Efficiency Varies Among Neuron Types in Vertebrates and Invertebrates. PLoS Computational Biology 6:e1000840.

Shannon CE (1948) A Mathematical Theory of Communication. Bell System Technical Journal 27:379–423.

Shen G, Zhao D, Dong Y, Li Y, Zeng Y (2023) Dive into the Power of Neuronal Heterogeneity. arXiv.

Shoval O, Sheftel H, Shinar G, Hart Y, Ramote O, Mayo A, Dekel E, Kavanagh K, Alon U (2012) Evolutionary trade-offs, pareto optimality, and the geometry of phenotype space. Science 336:1157–1160.

Sihn D, Kim SP (2019) A spike train distance robust to firing rate changes based on the earth mover’s distance. Frontiers in Computational Neuroscience 13:82.

Song S, Sjöström PJ, Reigl M, Nelson S, Chklovskii DB (2005) Highly Nonrandom Features of Synaptic Connectivity in Local Cortical Circuits. PLoS Biology 3:e68.

Stober TM, Batulin D, Triesch J, Narayanan R, Jedlicka P (2023) Degeneracy in epilepsy: multiple routes to hyperexcitable brain circuits and their repair. Communications Biology 6:479.

Tejero-Cantero A, Boelts J, Deistler M, Lueckmann JM, Durkan C, Gonçalves P, Greenberg D, Macke J (2020) sbi: A toolkit for simulation-based inference. Journal of Open Source Software 5:2505.

Tsodyks M, Pawelzik K, Markram H (1998) Neural Networks with Dynamic Synapses. Neural Computation 10:821–835.

Tsodyks MV, Markram H (1997) The neural code between neocortical pyramidal neurons depends on neurotransmitter release probability. Proceedings of the National Academy of Sciences 94:719–723.

Turrigiano G, Abbott LF, Marder E (1994) Activity-dependent changes in the intrinsic properties of cultured neurons. Science 264:974–977.

Turrigiano GG, Leslie KR, Desai NS, Rutherford LC, Nelson SB (1998) Activity-dependent scaling of quantal amplitude in neocortical neurons. Nature 391:892–896.

Turrigiano GG, Nelson SB (2004) Homeostatic plasticity in the developing nervous system. Nature Reviews Neuroscience 5:97–107.

Van Essen DC, Maunsell JH (1983) Hierarchical organization and functional streams in the visual cortex. Trends in Neurosciences 6:370–375.

Wahl-Schott C, Biel M (2008) HCN channels: Structure, cellular regulation and physiological function. Cellular and Molecular Life Sciences 66:470.

Wang Z, Ying Z, Bosy-Westphal A, Zhang J, Schautz B, Later W, Heymsfield SB, Müller MJ (2010) Specific metabolic rates of major organs and tissues across adulthood: evaluation by mechanistic model of resting energy expenditure. The American Journal of Clinical Nutrition 92:1369–1377.

Yang J, Prescott SA (2023) Homeostatic regulation of neuronal function: importance of degeneracy and pleiotropy. Frontiers in Cellular Neuroscience 17:1184563.

Yi GS, Wang J, Li HY, Wei XL, Deng B (2016) Metabolic Energy of Action Potentials Modulated by Spike Frequency Adaptation. Frontiers in Neuroscience 10:534.

Yu L, Shen Z, Wang C, Yu Y (2018) Efficient Coding and Energy Efficiency Are Promoted by Balanced Excitatory and Inhibitory Synaptic Currents in Neuronal Network. Frontiers in Cellular Neuroscience 12:123.

Yu L, Yu Y (2017) Energy efficient neural information processing in individual neurons and neuronal networks. Journal of Neuroscience Research 95:2253–2266.

Yu Y, Herman P, Rothman DL, Agarwal D, Hyder F (2017) Evaluating the gray and white matter energy budgets of human brain function. Journal of Cerebral Blood Flow & Metabolism 38:1339–1353.

Yuan Y, Huo H, Fang T (2018) Effects of Metabolic Energy on Synaptic Transmission and Dendritic Integration in Pyramidal Neurons. Frontiers in Computational Neuroscience 12:79.

Zeldenrust F, Calcini N, Yan X, Bijlsma A, Celikel T (2024) The tuning of tuning: How adaptation influences single cell information transfer. PLOS Computational Biology 20:e1012043.

Zeldenrust F, Gutkin B, Denéve S (2021) Efficient and robust coding in heterogeneous recurrent networks. PLoS Computational Biology 17:e1008673.

Zeldenrust F, Knecht Sd, Wadman WJ, Deneve S, Gutkin B (2017) Estimating the Information Extracted by a Single Spiking Neuron from a Continuous Input Time Series. Frontiers in Computational Neuroscience 11:49.

Zerlaut Y, Destexhe A (2017) A mean-field model for conductance-based networks of adaptive exponential integrate-and-fire neurons. arXiv.

Zhang M, Chitic R, Bohté SM (2025) Energy optimization induces predictive-coding properties in a multicompartment spiking neural network model. PLoS Computational Biology 21:e1013112.

Zhuchkova E, Remme MWH, Schreiber S (2013) Somatic versus Dendritic Resonance: Differential Filtering of Inputs through Non-Uniform Distributions of Active Conductances. PLoS ONE 8:e78908.

